# Comprehensive analysis of autophagic functions of WIPI family proteins and their implications for the pathogenesis of β-propeller associated neurodegeneration

**DOI:** 10.1101/2023.03.08.531694

**Authors:** Takahiro Shimizu, Norito Tamura, Taki Nishimura, Chieko Saito, Hayashi Yamamoto, Noboru Mizushima

## Abstract

β-propellers that bind polyphosphoinositides (PROPPINs) are an autophagy-related protein family conserved throughout eukaryotes. The PROPPIN family includes Atg18, Atg21, and Hsv2 in yeast and WD-repeat protein interacting with phosphoinositides (WIPI)1–4 in mammals. Mutations in the *WIPI* genes are associated with human neuronal diseases, including β-propeller associated neurodegeneration (BPAN) caused by mutations in *WDR45* (encoding WIPI4). In contrast to yeast PROPPINs, the functions of mammalian WIPI1–WIPI4 have not been systematically investigated. Although the involvement of WIPI2 in autophagy has been clearly shown, the functions of WIPI1, WIPI3, and WIPI4 in autophagy remain poorly understood. In this study, we comprehensively analyzed the roles of WIPI proteins by using *WIPI*-knockout (single, double, and quadruple knockout) HEK293T cells and recently developed HaloTag-based reporters, which enable us to monitor autophagic flux sensitively and quantitatively. We found that WIPI2 was nearly essential for autophagy and partially redundant with WIPI1. Autophagic flux was unaffected or only slightly reduced by single deletion of WIPI3 (encoded by *WDR45B*) or WIPI4 but was profoundly reduced by double deletion of WIPI3 and WIPI4. Furthermore, we revealed variable effects of BPAN-related missense mutations on the autophagic activity of WIPI4. BPAN is characterized by neurodevelopmental and neurodegenerative abnormalities, and we found a possible association between the magnitude of the defect of the autophagic activity of WIPI4 mutants and the severity of neurodevelopmental symptoms. However, some of the BPAN-related missense mutations, which produce neurodegenerative signs, showed almost normal autophagic activity, suggesting that non-autophagic functions of WIPI4 may be related to neurodegeneration in BPAN.

## Introduction

Macroautophagy (hereinafter, autophagy) is a degradation pathway that functions to maintain cellular homeostasis and is highly conserved among eukaryotes. In autophagy, a flat membrane cisterna termed the phagophore (also called the isolation membrane) expands to engulf cytoplasmic material, including proteins and organelles, and forms a double-membrane spherical structure called the autophagosome. Engulfed material is degraded after subsequent fusion with lysosomes. This pathway requires the concerted and hierarchical activities of autophagy-related (ATG) proteins (1–3). Upon starvation, the Atg1/ULK complex activates phosphatidylinositol 3-kinase complex 1, which generates phosphatidylinositol 3-phosphate (PI3P) in the endoplasmic reticulum (ER) and phagophore membranes. β-propellers that bind polyphosphoinositides (PROPPINs) are recruited to PI3P-rich membranes, further recruiting downstream factors, namely Atg2/ATG2 and the Atg12/ATG12–Atg5/ATG5–Atg16/ATG16L1 complex (1, 2). Atg2/ATG2 transfers phospholipids from the ER to phagophores to expand the membrane (4–6). Atg8/ATG8 family proteins (i.e., LC3 and GABARAP family proteins) are conjugated to phosphatidylethanolamine (PE) in autophagosomal membranes in a manner dependent on the Atg12/ATG12–Atg5/ATG5–Atg16/ATG16L1 complex (7–9).

The PROPPIN family includes Atg18, Atg21, and Hsv2 in yeast, Atg-18 and Epg-6 in *Caenorhabditis elegans*, and WD-repeat protein interacting with phosphoinositides (WIPI)1–4 in mammals. PROPPINs are phylogenetically divided into two paralog groups (10, 11): one group includes Atg18, Atg21, Atg-18, and WIPI1/2, and the other includes Hsv2, Epg-6, and WIPI3/4 (10, 11). The functions and hierarchical relationships of the three yeast PROPPINs have been well documented. Among them, Atg18 is essential for autophagy (12, 13); Atg18 forms a complex with Atg2, and the formation of this complex is necessary for the recruitment of both proteins to phagophores (14, 15). Atg21 is only partially required for autophagy; Atg21 interacts with Atg16 and recruits the Atg12–Atg5–Atg16 complex to induce Atg8–PE conjugation (16–18). Although Hsv2 is dispensable for autophagy, Hsv2 also interacts with Atg2 (19). In addition to autophagic functions, yeast PROPPINs have non-autophagic functions. For example, Atg18 is involved in the regulation of vacuolar size (20), vacuolar membrane fission (21), and transport from endo-lysosomal compartments to the Golgi apparatus (20,22,23). This function of Atg18 requires interaction with the retromer proteins Vps26 and Vps35, but this Atg18 complex is distinct from the retromer (22, 23). It has also been reported that Hsv2 is localized to endosomes under basal conditions, although its physiological role is unknown (19). In *C. elegans*, Epg-6 associates with Atg-2 and regulates autophagosome maturation, whereas Atg-18 functions in an earlier step, namely, autophagosome formation (24). Thus, the functions of the two paralog groups in autophagy, especially in the recruitment of Atg2 to phagophores, are complicated and even differ among species.

In contrast to yeast PROPPINs, the function of each WIPI protein and the functional relationships among WIPI1–4 have not been fully elucidated. Although WIPI2 has been shown to be involved in autophagy through recruiting ATG16L1 to phagophores (25), the roles of WIPI1, WIPI3, and WIPI4 in autophagy remain poorly understood, and it is still unclear whether WIPI proteins contribute to the recruitment of ATG2A/B to phagophores in mammalian cells. One study on the functions of WIPI1–4 showed that the stable knockdown of any WIPI impaired autophagosome formation in the human melanoma cell line G361 and suggested that WIPI3 and WIPI4 function downstream of WIPI1 and WIPI2 (26). Biochemical and structural studies revealed that WIPI4 strongly interacts with ATG2A/B (19,26,27), and other studies showed that ATG2A/B also interacts with WIPI3 to a lesser extent *in vivo* (28) and that a short peptide derived from ATG2A/B interacts with WIPI3 *in vitro* (29), suggesting possible contributions of WIPI3 and WIPI4 to the recruitment of ATG2A/B. Although WIPI3, WIPI4, and ATG2A/B are expressed ubiquitously, simultaneous deletion of WIPI3 and WIPI4 causes an autophagy defect observed only in cultured neuronal cells and mouse brains, but not in cultured non-neuronal cells, suggesting that WIPI3 and WIPI4 are important only in neuronal tissues (30, 31). Moreover, it was shown that an ATG2A mutant (residues Y1395, F1396, and S1397 all mutated into alanines), which is unable to interact with WIPI4, supported the degradation of p62 and LC3B during starvation (32), suggesting the interaction between ATG2A/B and WIPI4 (and likely WIPI3) (28, 29) is dispensable for autophagic flux. It was also shown that ATG2A/B proteins are recruited to phagophores by the interaction between ATG2A/B and ATG8 family proteins such as GABARAP, GABARAPL1, and LC3A (32). Therefore, the functional relationships among WIPI proteins seem to be complicated and merit further clarification.

Mutations in *WIPI* genes are known to cause human neuronal diseases. For example, mutations in *WDR45*, which encodes WIPI4, cause β-propeller protein-associated neurodegeneration (BPAN, formally called static encephalopathy of childhood with neurodegeneration in adulthood [SENDA], OMIM#300894), which is typically characterized by static neurodevelopmental symptoms in the early stage of the disease and neurodegeneration in the late stage of the disease (33–35). Mutations in *WDR45B*, which encodes WIPI3, also cause a complex neurodevelopmental disorder called El-Hattab-Alkuraya syndrome (OMIM#617977) (36–38), while mutations in *WIPI2* cause a neurodevelopmental disorder accompanied by skeletal and cardiac abnormalities (OMIM#618453) (39, 40). To understand the pathogenesis of these WIPI-related diseases, it would be helpful to reveal the function of each WIPI protein more precisely.

Because the contributions of each WIPI protein and of the interaction of ATG2 with WIPIs and ATG8s to autophagy may be only partial, more sensitive, quantitative methods for monitoring autophagic flux would be necessary for a systematic analysis of these proteins. Recently, we developed a highly sensitive and quantitative method, the HaloTag (Halo)-based reporter processing assay (41). Using this method, we comprehensively analyzed the function of individual WIPI proteins using single or multiple *WIPI*-knockout HEK293T cells and revealed the necessity of WIPI2 in autophagy and the rigorous contribution of WIPI4 to the recruitment of ATG2 to phagophores. Finally, we found a possible association between the magnitude of autophagy defect and the severity of neurodevelopmental symptoms by assessing the autophagic activities of disease-related mutants of WIPI3 and WIPI4. In contrast, no clear correlation was found between the magnitude of autophagy defects and the age of the onset of neurodegeneration, suggesting that neurodegeneration in BPAN may be caused by a defect in a non-autophagic function of WIPI4.

## Results

### WIPI2 is nearly essential for mammalian autophagy

To investigate the importance of individual WIPI proteins in mammalian autophagy, we generated *WIPI1–4* single knockout (KO) and *WIPI1–4* quadruple KO (QKO) HEK293T cells (Fig. S1). In addition, because of the close phylogenetic relationships between WIPI1 and WIPI2 and between WIPI3 and WIPI4 (10, 11), we generated *WIPI1/2* double KO (DKO) and *WIPI3/4* DKO HEK293T cells (Fig. S1). As ATG16L1 and ATG2A/B function immediately downstream of WIPIs (1, 2), we also generated *ATG16L1* KO and ATG2A/B DKO HEK293T cells (Fig. S1). To measure the autophagic activity of these cells using the Halo-based reporter processing assay, we stably introduced the Halo–monomeric GFP (Halo–GFP) and Halo–LC3B (simply referred to as Halo–LC3) reporters (41). When these Halo-based reporters are delivered to lysosomes via autophagy, they are efficiently degraded. However, Halo becomes resistant to lysosomal degradation when it is conjugated with Halo ligands, producing a free Halo fragment in an autophagy-dependent manner (Fig. 1A).

**Figure 1.**
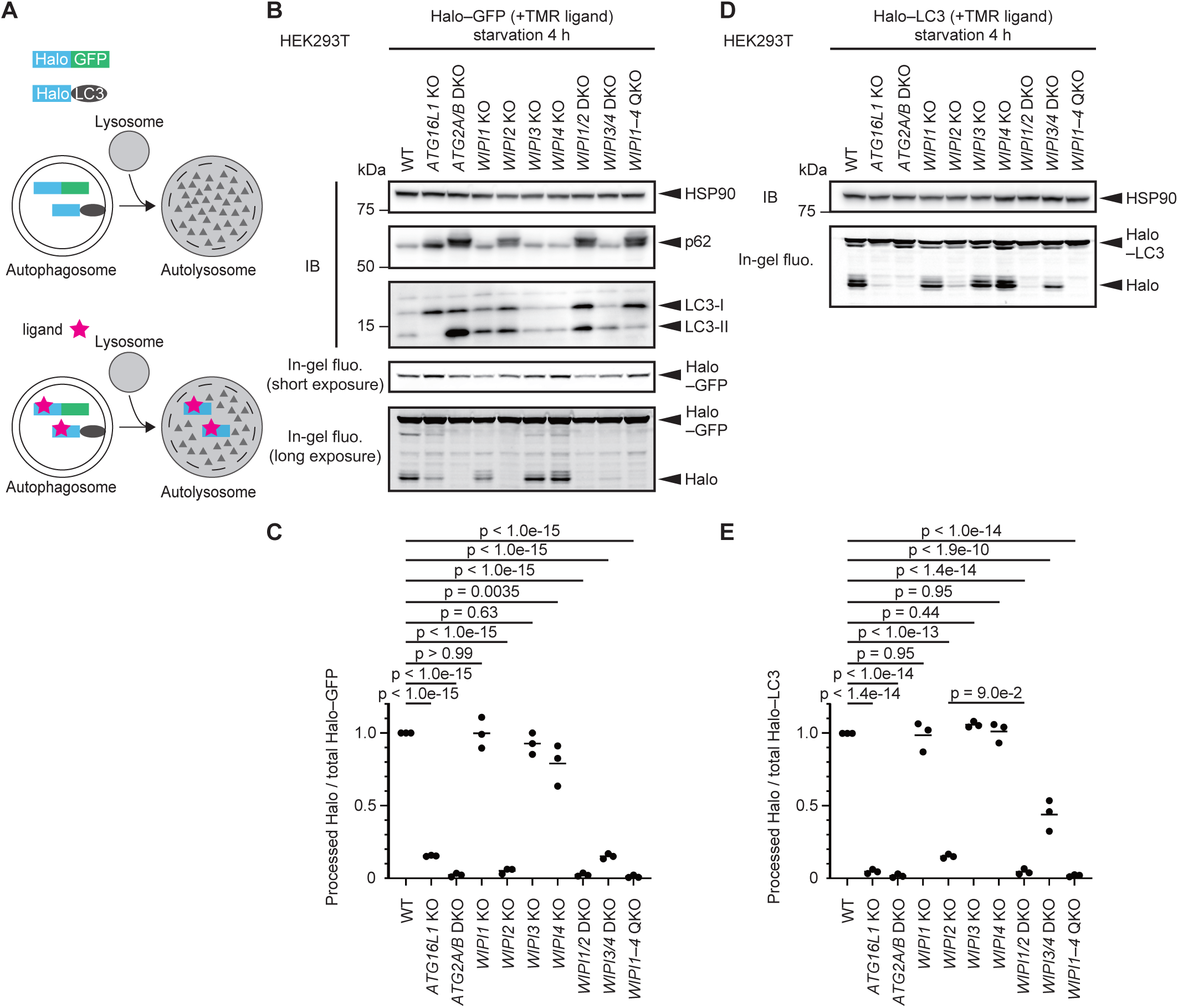
Analysis of autophagic flux in *WIPI* KO cells. (A) Schematic representation of the Halo processing assay. Halo–GFP and Halo–LC3 are incorporated into autophagosomes non-selectively and selectively, respectively. Without ligand binding, these reporters are degraded in autolysosomes. In contrast, after ligand binding, Halo becomes stable and accumulates in autolysosomes, the amount of which reflects autophagic flux. (B and D) Halo–GFP and Halo–LC3 processing assay. Cells stably expressing Halo–GFP (B) or Halo–LC3 (D) were labeled for 30 min with 100 nM tetramethylrhodamine (TMR)-conjugated Halo ligand and then were incubated in starvation medium for 4 h. Total cell lysates were subjected to immunoblotting with the indicated antibodies or in-gel fluorescence detection. WT, wild-type. (C and E) Results from (B) and (D) were quantified. The Halo–GFP/LC3 processing rate was calculated as the band intensity of processed Halo over that of Halo–GFP/LC3.The plots show the Halo–GFP/LC3 processing rate of each cell line relative to that in WT cells. Solid bars indicate the means, and dots indicate the data from three independent experiments. Data were statistically analyzed by one-way ANOVA with Dunnett’s test (C) or the Holm-Šídák test (E).

First, we used the Halo–GFP reporter, which is incorporated into autophagosomes non-selectively. When wild-type HEK293T cells stably expressing Halo–GFP were incubated with a tetramethylrhodamine (TMR)-conjugated ligand for 30 min and further incubated in a ligand-free amino-acid starvation medium for 4 h, the band corresponding to free Halo was observed by in-gel fluorescence detection (Fig. 1B). In *ATG16L1* KO cells, Halo–GFP processing was markedly impaired, but a faint Halo fragment was still detected, which is consistent with previous findings that a low level of autophagic flux remains in cells lacking the ATG8 conjugation machinery (42–44). Halo–GFP processing was completely blocked in *ATG2A/B* DKO cells.

Among *WIPI* single KO cells, *WIPI2* KO cells showed an almost complete absence of Halo–GFP processing, suggesting that WIPI2 is nearly essential for autophagy (Fig. 1B and C). *WIPI1* KO and *WIPI3* KO had no significant impact on Halo–GFP processing, whereas *WIPI4* KO caused a slight reduction. Halo–GFP processing was further impaired in *WIPI3/4* DKO cells, indicating the functional redundancy of WIPI3 and WIPI4, as previously observed in neuronal cells (30).

Next, we conducted the processing assay using the Halo–LC3 reporter, which is selectively sequestered into autophagosomes. Again, the processing of Halo–LC3 was completely blocked in both *ATG2A/B* DKO and *WIPI1–4* QKO cells (Fig. 1D and E).

*WIPI2* KO cells showed an almost complete absence of Halo–LC3 processing, and the residual Halo–LC3 processing was further impaired by the simultaneous knockout of *WIPI1* and *WIPI2* (in *WIPI1/2* DKO cells), suggesting that, although WIPI2 plays a dominant role, WIPI1 functions redundantly with WIPI2 (Fig. 1D and E). *WIPI3/4* DKO cells showed a partial but still significant impairment of Halo–LC3 processing. Unlike the Halo–GFP processing assay, the Halo–LC3 processing assay showed no significant defect in *WIPI4* KO cells. This may be caused by a difference in the dynamic range or sensitivity between the Halo–GFP and Halo–LC3 processing assays. Because Halo–LC3 is selectively and efficiently incorporated into autophagosomes, the Halo–LC3 processing assay has higher sensitivity to detect low autophagic activities (e.g., in *WIPI2* KO cells). However, when autophagic activities are relatively high (e.g., in *WIPI4* KO cells), Halo–LC3 processing can become easily saturated and accordingly unable to reveal a subtle decrease in autophagic flux. Thus, the Halo–GFP processing assay would be more suitable for detecting a slight defect.

In contrast to our Halo-based processing assay showing a clear decrease in autophagic flux in *WIPI3/4* DKO HEK293T cells (Fig. 1), a recent study suggested that WIPI3 and WIPI4 are dispensable for autophagy in non-neuronal cells, as p62 (also known as SQSTM1) and LC3 do not accumulate in *WIPI3/4* DKO COS7 cells and mouse embryonic fibroblasts (30). Consistently, we confirmed no accumulation of lipidated LC3 (LC3-II) and p62 as well as normal degradation of LC3-II during starvation in *WIPI3/4* DKO HEK293T cells (Fig. 1B, S2A–C). We also assessed autophagic flux in *WIPI3/4* DKO HEK293T cells using another reporter, GFP–LC3–RFP (Fig. S2D) (42). GFP–LC3–RFP is cleaved by endogenous ATG4 proteins to produce an equimolar amount of GFP–LC3 and RFP. GFP–LC3 is conjugated to autophagosomal membranes and then degraded or quenched, while RFP remains in the cytosol. Therefore, the reduction in the GFP:RFP fluorescence ratio represents autophagic flux. We observed that the GFP:RFP fluorescence ratio was similar in wild-type and *WIPI3/4* DKO HEK293T cells before and after starvation (Fig. S2E). Thus, the conventional methods using LC3 and p62 were unable to detect the slight autophagy defect in *WIPI3/4* DKO HEK293T cells. This is probably due to the lower sensitivity of these degradation-based assays, which measure the reduction of autophagy substrates, compared with the Halo-based processing assay, which measures the production of the Halo fragment (i.e., positive signal).

In summary, in HEK293T cells, WIPI2 is nearly essential for autophagy, and WIPI1 functions redundantly with WIPI2, although its contribution is limited. WIPI3 and WIPI4 function redundantly for autophagy, and WIPI3 and WIPI4 are each only partially necessary or not necessary for autophagy. Considering the non-selective incorporation of Halo–GFP into autophagosomes, the Halo–GFP processing assay should reflect the true activity of bulk autophagy. Therefore, we used it in the following experiments.

### WIPI2 and WIPI4 regulate autophagy at different steps

Next, we introduced individual WIPIs or combinations of WIPIs into *WIPI* QKO cells. Among two WIPI1 isoforms (A and B) and six WIPI2 isoforms (A, B, C, D, E, and delta), WIPI1A, WIPI2B, and WIPI2D have been reported to be recruited to phagophores (10,11,25,26,43). In particular, WIPI2B has been shown to play an important role in autophagy *in vivo* (25). Therefore, we used WIPI1A and WIPI2B for this experiment. When expressed alone, only WIPI2B restored Halo–GFP processing in *WIPI* QKO cells (Fig. 2A and B). The p62 level was also reduced. The restoration of Halo–GFP processing by WIPI2 was not changed by the further addition of WIPI1A or WIPI3, but was significantly enhanced by the addition of WIPI4 (Fig. 2A and B). Almost full recovery was observed when both WIPI2B and WIPI4 were expressed. WIPI3 and WIPI4 could not restore Halo–GFP processing, even when both proteins were expressed (Fig. 2A and B). These results suggest that WIPI2 and WIPI4 support autophagy at different steps and that WIPI2 is essential for the function of WIPI4 in autophagy.

**Figure 2.**
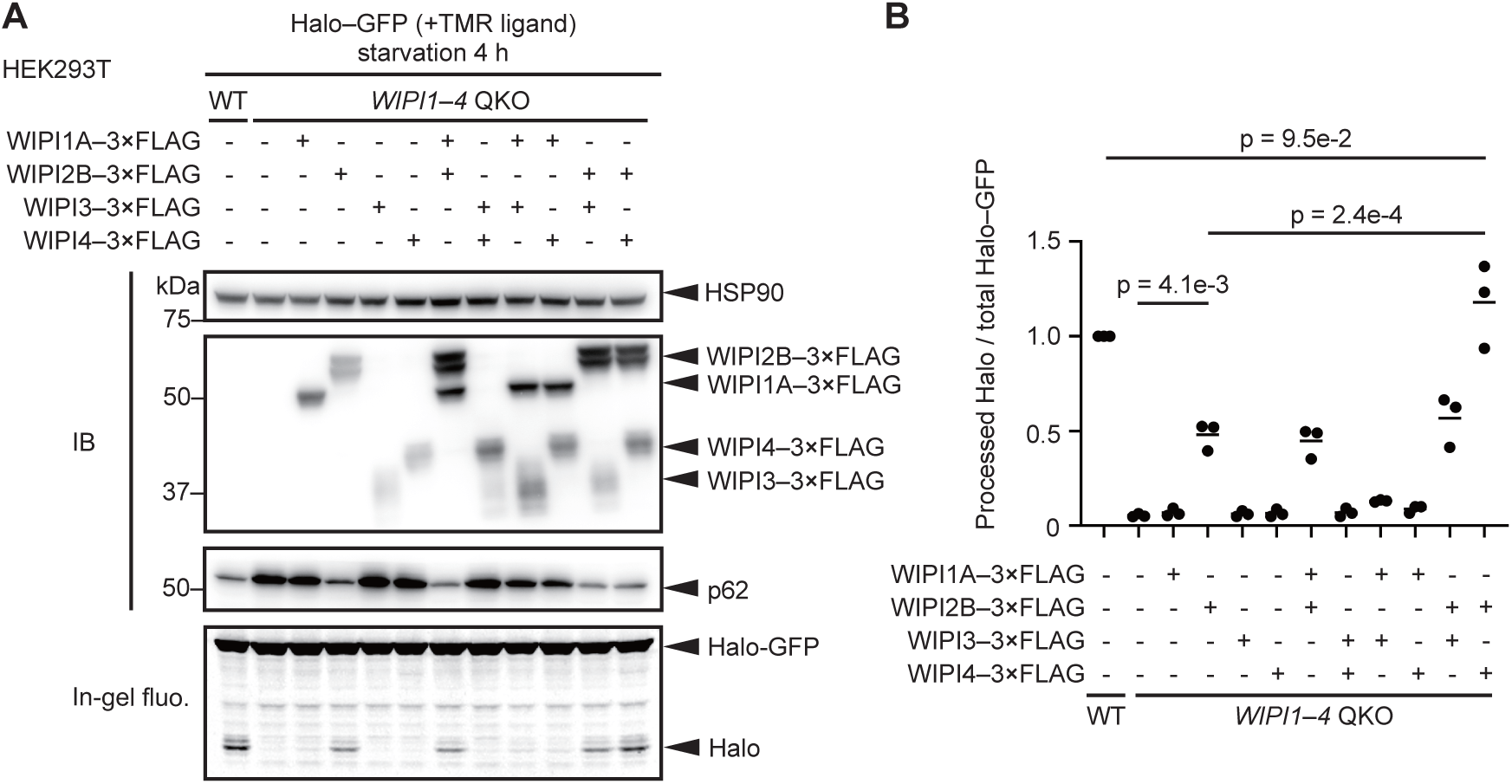
WIPI2 and WIPI4 regulate autophagy through different mechanisms. (A) Halo–GFP processing assay of WT and WIPI1–4 QKO HEK293T cells with or without indicated 3×FLAG-tagged WIPI proteins was performed as in Figure 1B. (B) Data from (A) were quantified, and the results are shown as described in Figure 1. Solid bars indicate the means, and dots indicate the data from three independent experiments. Data were statistically analyzed by one-way ANOVA with the Holm-Šídák test.

### WIPI4 regulates autophagy through its interaction with PI3P and ATG2A/B and maintains the size of autophagosomes

By using Halo-based reporters, we were able to detect a clear autophagy defect in *WIPI3/4* DKO cells, which was difficult to detect by using conventional methods (Fig. 1, S2), enabling us to further examine the roles of WIPI3 and WIPI4 in autophagy. To investigate the significance of PI3P binding of WIPI3 and WIPI4, we generated WIPI3 R225T/R226T (WIPI3 LTTG) and WIPI4 R232T/R233T (WIPI4 LTTG) mutants, which completely lost their binding to PI3P (26,44,45). In *WIPI3/4* DKO cells, the expression of either WIPI3 or WIPI4 significantly restored Halo–GFP processing, confirming that WIPI3 and WIPI4 are indeed functionally redundant (Fig. 3A and B). In contrast, the WIPI3 LTTG and WIPI4 LTTG mutants failed to restore Halo–GFP processing, suggesting that WIPI3 and WIPI4 function through binding to PI3P (Fig. 3A and B).

**Figure 3.**
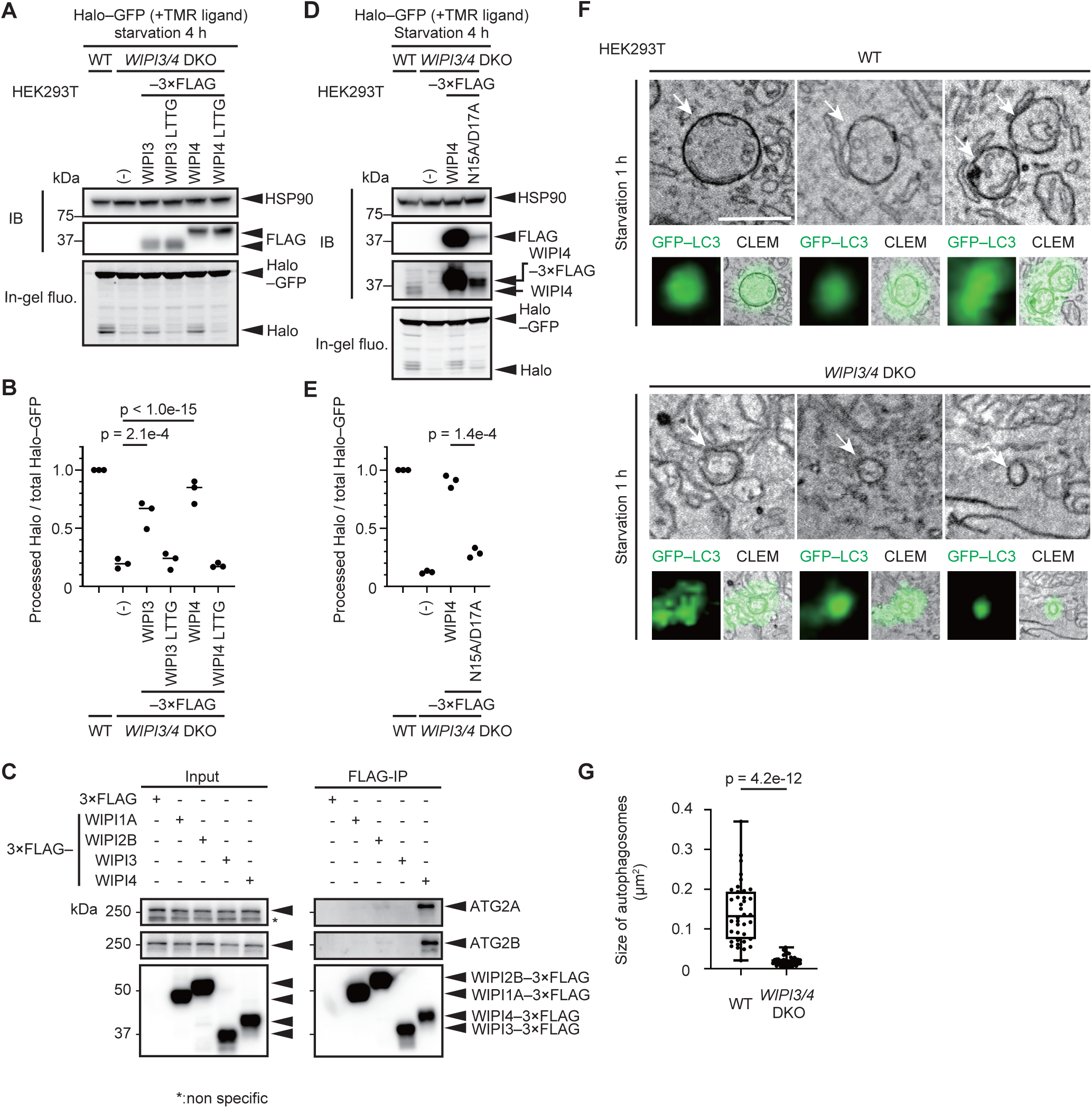
WIPI3 and WIPI4 regulate autophagy through their interactions with PI3P and ATG2 and maintain the size of autophagosomes. (A) and (D) Halo–GFP processing assay of WT cells and *WIPI3/4* DKO HEK293T cells expressing 3×FLAG-tagged WIPI3/4 (as indicated) and their mutants unable to bind PI3P or ATG2 was performed as in Figure 1B. (B) and (E) Data from (A) and (D) were quantified and the results are shown as described in Figure. 1. Solid bars indicate the means, and dots indicate the data from three independent experiments. Data were statistically analyzed by Dunnett’s test (B) and Welch’s *t*-test (E). (C) Cell lysates from HEK293T cells transiently expressing FLAG-tagged WIPI1–4 were subjected to immunoprecipitation with anti-FLAG antibody and immunoblotting with antibodies against FLAG and endogenous ATG2A and ATG2B. (F) Representative autophagosomes (arrows) of WT and *WIPI3/4* DKO HEK293T cells identified in 3D correlative light and electron microscopy (3D-CLEM) images using GFP–LC3 as a marker. Scale bar, 500 nm. (G) Maximum cross-sectional areas of autophagosomes identified by 3D-CLEM were quantified. Data were collected from 37 autophagosomes in three WT cells and 42 autophagosomes in three *WIPI3/4* DKO cells. Data were statistically analyzed by Welch’s *t*-test.

WIPI4 is known to interact with ATG2A/B (26,28,29). We confirmed that ATG2A/B interacts with WIPI4 almost exclusively among WIPI1–4 (Fig. 3C). WIPI2B appeared to faintly interact with ATG2A/B. WIPI1A and, in contrast to the results of previous studies (28, 29), WIPI3 did not interact with ATG2A/B in this experimental condition. Therefore, we investigated whether the WIPI4–ATG2A/B interaction is necessary for the function of WIPI4. For this purpose, the WIPI4 N15A/D17A mutant, which is unable to interact with ATG2A/B, was prepared (26) (Fig. S3). Halo–GFP processing was significantly impaired in WIPI4 N15A/D17A-expressing *WIPI3/4* DKO cells in contrast to almost normal processing in wild-type WIPI4-expressing cells (Fig. 3D and E). In this experiment, the expression level of WIPI4 N15A/D17A was remarkably decreased compared with wild-type WIPI4, but the level of WIPI4 N15A/D17A was still higher than that of endogenous WIPI4 (Fig. 3D). These results suggest that the interaction between WIPI4 and ATG2 is important for autophagy. (28, 29)

It was previously reported that the size of autophagosomes is smaller in mouse *WIPI3/4* DKO Neuro-2a cells compared with parental cells (30). To confirm this pattern in HEK293T cells, we observed autophagosomes using 3D correlative light electron microscopy using GFP–LC3 as a marker (46). After a 1-h starvation period, GFP–LC3-positive closed autophagosomes were abundant in both wild-type and *WIPI3/4* DKO cells (Fig. 3F, S4A). Furthermore, we confirmed that the number of GFP–LC3-positive structures was not reduced in *WIPI3/4* DKO cells (Fig. S4B). However, the size of autophagosomes was strikingly reduced in *WIPI3/4* DKO cells (Fig. 3F and G). Collectively, these results suggest that WIPI4, and possibly WIPI3 also, support autophagic flux through interactions with PI3P and ATG2A/B and maintain the size, rather than the number, of autophagosomes even in non-neuronal cells.

### Recruitment of ATG2A to phagophores is mediated by its interaction with WIPI4 and ATG8s

Regarding the recruitment of ATG2A/B to phagophores, a previous study showed the significance of their interaction with ATG8s and suggested that the interaction between WIPI3/4 and ATG2A/B is dispensable for autophagy (32). However, using the Halo–GFP reporter, we instead found that the interaction between WIPI4 and ATG2A/B is crucial (Fig. 3D and E). Therefore, we next investigated the relative contributions of these two interactions using Halo-based reporters. For this purpose, we prepared several mutants for ATG2A only, as ATG2A and ATG2B function redundantly (47) (Fig. 4A). ATG2A interacts with WIPI3/4 through its WIPI-interacting region (WIR, residues 1374–1404) (29), while ATG2A interacts with ATG8s through its LC3-interacting region (LIR, residues 1362–1365) (32). We mutated these regions to render ATG2A unable to interact with either ATG8s or WIPI3/4 or both (Fig. 4A). We confirmed that LIR mutation reduced the interaction with GABARAP but not WIPI4, whereas WIR deletion almost completely blocked the interaction with WIPI4 but not GABARAP (Fig. S5A and B).

**Figure 4.**
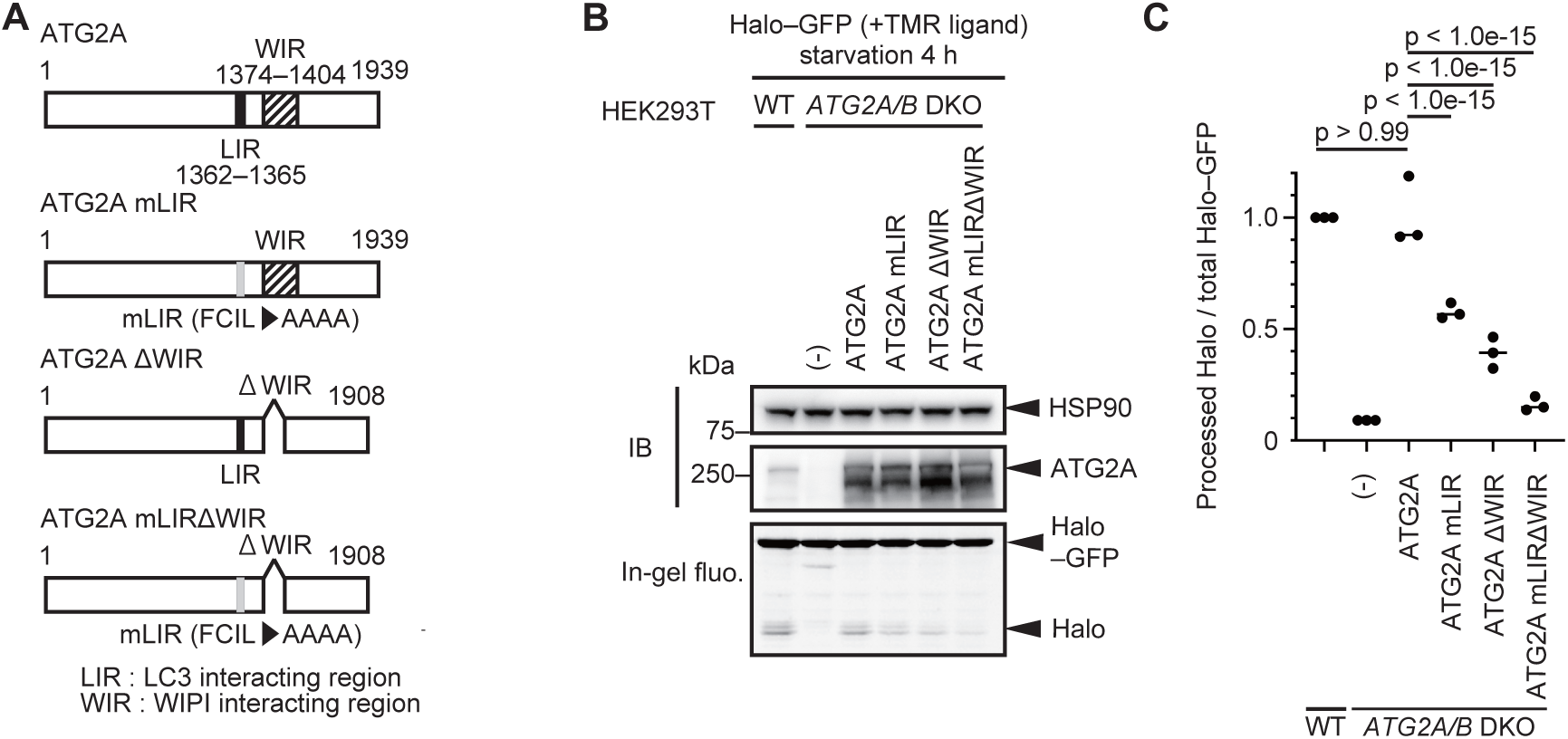
Interactions of ATG2A with both ATG8s and WIPI4 collaboratively contribute to the autophagic function of ATG2A. (A) Structures of ATG2A and its mutants. (B) The Halo–GFP processing assay of WT and *ATG2A/B* DKO HEK293T cells expressing ATG2A and its mutants was performed as in Figure 1B. (C) Data from (B) were quantified, and the results are shown as described in Figure 1C. Solid bars indicate the means, and dots indicate the data from three independent experiments. Data were statistically analyzed by one-way ANOVA with Dunnett’s test.

In *ATG2A/B* DKO cells, wild-type ATG2A fully rescued Halo–GFP processing (Fig. 4B and C). In contrast, the LIR mutant (ATG2A mLIR) or the WIR deletion mutant (ATG2A ΔWIR) showed partially impaired Halo–GFP processing (Fig. 4B and C). ATG2A with both LIR mutation and WIR deletion (ATG2A mLIRΔWIR) showed a further decrease in Halo–GFP processing, suggesting that LIR and WIR collaboratively function (Fig. 4B and C). These ATG2A mutants still interacted with WIPI2B to similar extents as wild-type ATG2A (Fig. S5C), suggesting that the interaction between ATG2A and WIPI2B is not important for autophagy.

Next, we investigated whether the reduced autophagic activity of the ATG2A mutants was attributable to defects in their recruitment to phagophores. ATG2A colocalized with WIPI2B, a phagophore marker, and LipidTOX, a lipid droplet marker, as previously reported (47, 48) (Fig. 5A and B). In contrast, the colocalization rates between ATG2A mutants (mLIR, ΔWIR, and mLIRΔWIR) and WIPI2B were significantly reduced (Fig. 5A and B). Instead, most puncta of the ATG2A mutants colocalized with LipidTOX. These results suggest that the interactions of ATG2A with WIPI4 and ATG8s contribute to the recruitment of ATG2A to phagophores.

**Figure 5.**
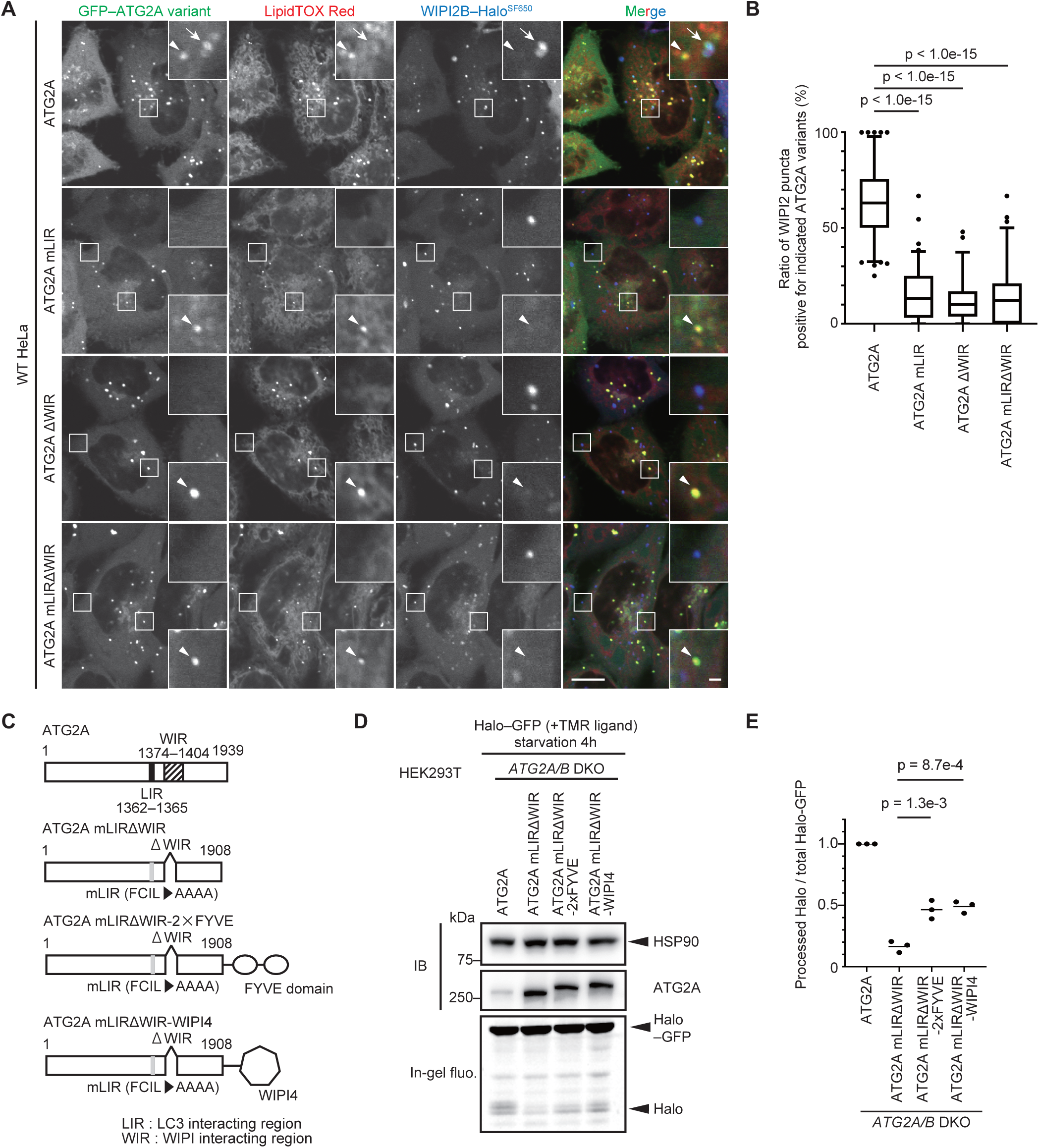
The recruitment of ATG2A to phagophores is mediated by its interaction with WIPI4 and ATG8s. (A) HeLa cells stably expressing muGFP–ATG2A or the indicated ATG2A mutants with WIPI2B–Halo were observed by confocal microscopy after being incubated in starvation medium with 200 nM SF650-conjugated HaloTag ligand and LipidTOX Red for 1 h. Arrows indicate the colocalization of a muGFP–ATG2A punctum with a WIPI2B–Halo^SF650^ punctum. Arrowheads indicate the colocalization of puncta of muGFP–ATG2A or ATG2A mutants with LipidTOX Red puncta. Scale bars indicate 10 μm (main image) and 1 μm (inset). (B) The ratio of WIPI2B–Halo^SF650^ puncta colocalizing with muGFP puncta was calculated. Data were collected from more than 90 cells for ATG2A or ATG2A mutants. The box plot denotes the median and 2.5–97.5 percentiles, while outliers are plotted individually. Data were statistically analyzed by one-way ANOVA with Dunnett’s test. (C) Structures of ATG2A, the mLIRΔWIR mutant, and ATG2A mLIRΔWIR fused with the 2×FYVE domain or WIPI4. (D) The Halo–GFP processing assay of *ATG2A/B* DKO HEK293T cells expressing the indicated proteins was performed as in Figure 1B. (E) Data from (D) were quantified, and the results are shown as described in Figure 1C. Solid bars indicate the means, and dots indicate the data from three independent experiments. Data were statistically analyzed by one-way ANOVA with Dunnet’s test.

To further confirm that the defects of the ATG2A mutants were indeed attributable to impaired recruitment, we assessed whether the forced recruitment of ATG2A mLIRΔWIR to phagophores could recover the function of ATG2A. We fused the PI3P-binding 2×FYVE domain (49) or WIPI4 to ATG2A mLIRΔWIR (Fig. 5C). These fusion proteins significantly enhanced Halo–GFP processing in *ATG2A/B* DKO cells compared with ATG2A mLIRΔWIR, confirming that proper recruitment of ATG2A is important (Fig. 5D and E).

### Some of the disease-associated WIPI4 mutants retain autophagic activity

Next, we evaluated the autophagic activity of disease-related mutants of WIPI3 and WIPI4 (34,36,50–60). One WIPI3 mutant and 15 WIPI4 mutants were expressed in *WIPI3/4* DKO HEK293T cells, and Halo–GFP processing after a 4-h starvation period was evaluated. We used *WIPI3/4* DKO cells for this purpose because *WIPI3* KO and *WIPI4* KO HEK293T cells showed no or only slight autophagy defects (Fig. 1). The WIPI3 R109Q mutant showed decreased Halo–GFP processing (the clinical phenotype was not described in detail (36)) (Fig. 6A and B). The disease-related WIPI4 mutations exhibited various effects on Halo–GFP processing. Notably, four WIPI4 mutants (V66E, D84G, F100S, and N202K) showed almost normal Halo–GFP processing similar to that of wild-type WIPI4, and two mutants (R134P and G168E) showed only mild defects (Fig. 6A and B).

**Figure 6.**
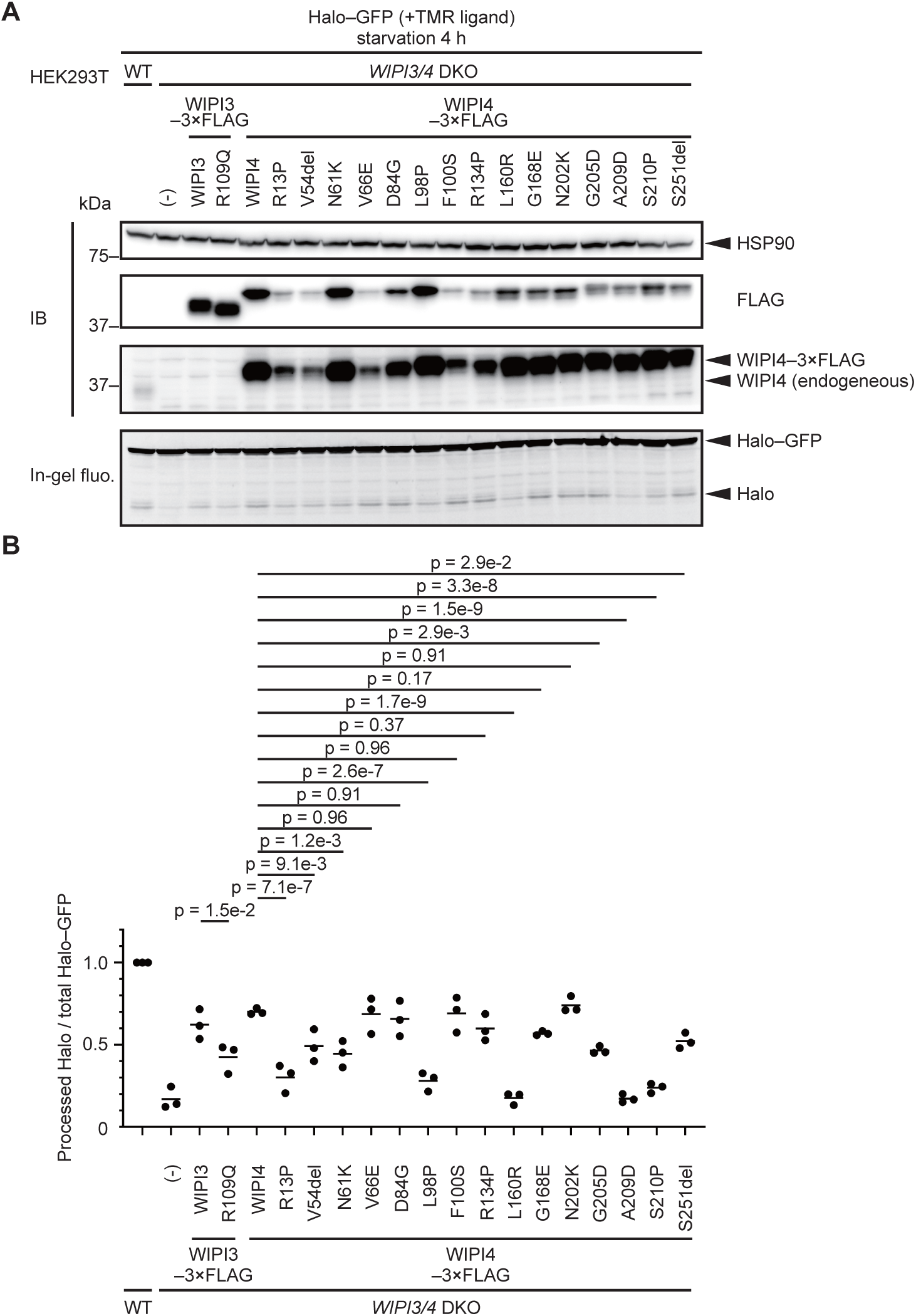
Some of the disease-associated WIPI4 mutants retain autophagic activity. (A) Halo–GFP processing assay of WT and *WIPI3/4* DKO HEK293T cells with or without 3×FLAG-tagged WIPI3/4 or their disease-related mutants was performed as in Figure 1B. (B) Data from (A) were quantified, and the results are shown as described in Figure 1C. Solid bars indicate the means, and dots indicate the data from three independent experiments. Data were statistically analyzed by one-way ANOVA with the Holm-Šídák test.

Furthermore, we investigated the relationship between the magnitude of autophagy defect and the clinical phenotype of each mutation in WIPI4. Patients with these mutations vary in age, and some of the patients may have not yet developed symptoms indicative of neurodegeneration, such as parkinsonism and dementia. In addition, clinical phenotypes were not described in detail for all the patients. Therefore, we first focused on the severity of intellectual disability, which is relatively detailed in most of the reports, as an indicator of the severity of neurodevelopmental symptoms. We examined 2 male patients and 14 female patients (34,50–60). In 5 female patients with mutations D84G, F100S, R134P, G168E, and N202K, which were determined to have no impact or only a mild impact on autophagic activity, intellectual disability tended to be mild (52–54,56). Patients with mutations D84G and F100S could speak in sentences and began to walk at a typical age (52, 53). Patients with mutations R134P and G168E, who were 2-year-7-month-old and 5-year-old patients, respectively, could each speak a couple of words (54). The patient with mutation N202K could not communicate verbally but could communicate partially in a non-verbal way (56). In contrast, the intellectual disabilities of patients with mutations L160R, A209D, and S210P, which caused a profound defect in autophagic activity, seemed to be more severe, as they did not acquire any language abilities (55,58,59). Because *WDR45*, the gene encoding WIPI4, is on the X chromosome, the expression level of WIPI4 depends on the extent of X chromosome inactivation in each female case. In addition, the stabilities of mutant proteins may affect the expression level of each mutant WIPI4. Nevertheless, our data suggest a possible association between autophagic activity and the severity of intellectual disability.

Seven female patients developed symptoms indicative of neurodegeneration, such as dementia and parkinsonism (35,56,58–60). However, there was no clear correlation between the autophagic activity and the age of onset of these symptoms (Fig. 7). For example, although the N202K mutation had almost no impact on the autophagic activity of WIPI4, the patient with the N202K mutation developed parkinsonism at the age of 27 (56). Additionally, a patient with the D84G mutation, which also had no impact on the autophagic activity of WIPI4, showed iron accumulation in both the globus pallidus and substantia nigra, as detected by brain magnetic resonance imaging, indicating that the patient would likely develop parkinsonism in the future. Neurodevelopmental symptoms were very mild in this patient, probably reflecting almost normal autophagic flux. These findings suggest that WIPI4 has non-autophagic functions, which may be important for the prevention of neurodegeneration and iron accumulation.

**Figure 7.**
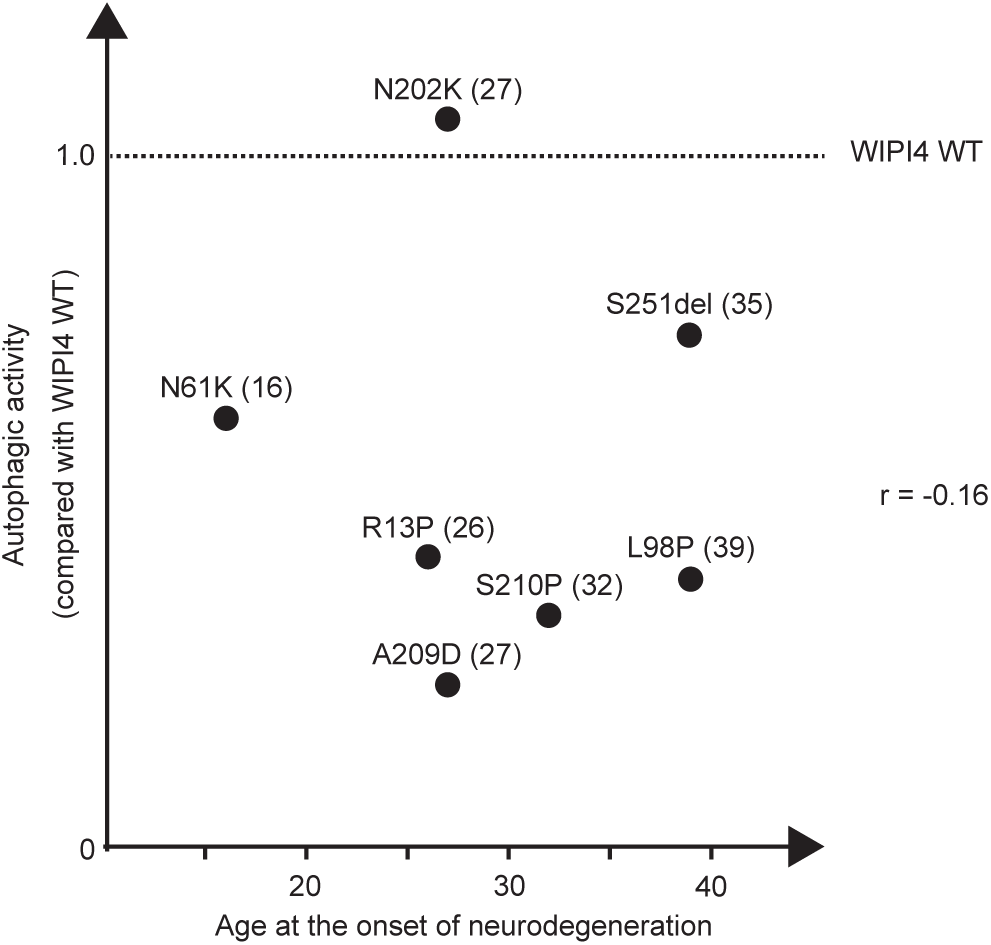
The magnitude of autophagy defects in WIPI4 mutants is possibly associated with the severity of neurodevelopmental symptoms but not with the age of neurodegeneration onset. The autophagic activity (%WT in Figure 6B) of corresponding WIPI4 mutants and the age of the onset of neurodegeneration symptoms, such as parkinsonism and dementia, were plotted for seven BPAN cases. The parenthetical after each mutation indicates the age of the onset of these symptoms. The Pearson’s correlation coefficient (*r*) between the degree of autophagic defect and the age of the onset of these symptoms is shown.

## Discussion

In this study, we conducted a systematic analysis of WIPI proteins using highly sensitive autophagic flux reporters and multiple knockout cell lines. We confirmed the nearly essential function of WIPI2 in autophagy (Fig. 1), which is consistent with a previous report showing that the depletion of WIPI2 caused the most drastic inhibition of autophagosome formation among the depletions of each of the WIPI proteins (26). It was also previously shown that the depletion of WIPI1 moderately inhibits autophagosome formation in G361 cells (26). However, our quantitative study in HEK293T cells showed, although WIPI1 may be redundant with WIPI2 possibly through its interaction with ATG16L1 (25, 26), the contribution of WIPI1 was very limited (Fig. 8). This result may correspond to the finding that the interaction between WIPI1 and ATG16L1 is far weaker than that between WIPI2 and ATG16L1 (25). Instead, WIPI1 is endowed with a non-autophagic function related to endosomal membrane fission and endosomal transport (61), which may correspond to the non-autophagic function of yeast Atg18 in retrograde transport from endosomes to the Golgi apparatus (20,22,23) (Fig.8) .

**Figure 8.**
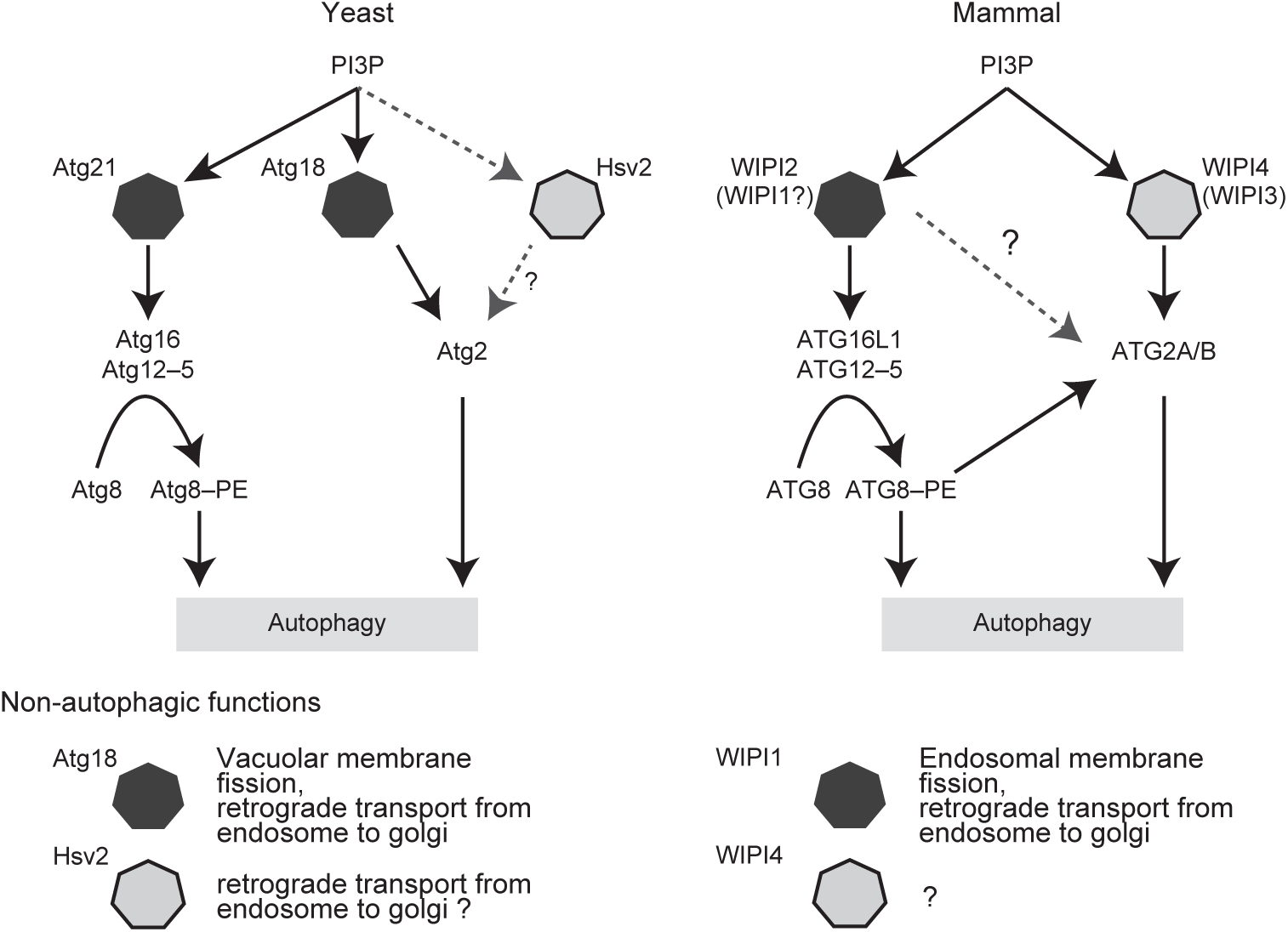
Proposed functions of PROPPINs. In yeast, Atg21 recruits Atg16, and Atg18 and possibly Hsv2 recruit Atg2. Atg18 is essential, and Atg21 is partially essential for autophagy. Hsv2 is dispensable for autophagy. In mammals, WIPI2 is predominantly important for autophagy, at least through binding with ATG16L1. WIPI1 may be partially redundant with this function of WIPI2. WIPI4 and possibly WIPI3 are important for the recruitment of ATG2A/B to the site of autophagosome formation. ATG2A/B recruitment is partially achieved through binding with ATG8. Phylogenetically related proteins are shown in the same colors (black or gray).

The roles of WIPI3 and WIPI4 in autophagy have been controversial. LC3 accumulates upon double knockdown of WIPI3 and WIPI4 in G361 cells (26). Although p62 accumulates in the brain (but not in the liver or kidney) of *Wipi3/4* DKO mice, it is virtually absent from the brain of *Wipi3* KO and *Wipi4* KO mice (31). The degradation of LC3 is reduced in fibroblasts derived from BPAN patients (62). On the other hand, it was reported that the levels of LC3 and p62 were normal in *WIPI3/4* DKO COS7 cells and *Wipi3/4* DKO mouse embryonic fibroblasts (30). In the present study, our Halo-based processing assay clearly revealed a decrease in autophagic flux in *WIPI3/4* DKO HEK293T cells, while canonical methods, such as LC3 and p62 degradation and GFP–LC3–RFP reporter assays, failed to detect the autophagy defect in *WIPI3/4* DKO HEK293T cells. We assume that these previous inconsistencies might be a consequence of the relatively low sensitivities of LC3-based autophagic flux assays to detect subtle autophagy defects. Our results using appropriate methods showed that WIPI3 and WIPI4 are redundantly important for autophagy, even in non-neuronal cells. Our data also suggest that, at least for autophagic flux in HEK293T cells, WIPI4 appears to be more important than WIPI3, because the deletion of WIPI4 showed a larger suppressive effect on autophagy than that of WIPI3 (Fig. 1), and the rescue effect of re-expression of WIPI4 in *WIPI1–4* QKO was greater than that of WIPI3 (in the presence of WIPI2) (Fig. 2). This may be because ATG2 interacts with WIPI4 more strongly than with WIPI3 (28). ATG2 was detected in the interactome of WIPI4 but not WIPI3 (26). Consistent with these results, we detected interaction between WIPI4 and ATG2A/B, but not between WIPI3 and ATG2A/B (Fig. 3C). Nevertheless, our results showing functional redundancy between WIPI3 and WIPI4 suggest that WIPI3 and WIPI4 function in autophagy through the same mechanism or at least a partially shared mechanism.

The next issue regards the relationship between the two functional units, WIPI1/2 and WIPI3/4. It was reported that, in *C. elegans*, Atg-18 (a homolog of WIPI1/2) appears to function upstream of Epg-6 (a homolog of WIPI3/4), but the phenotype of *atg-18* mutants is milder than that of *epg-6* mutants (24). In mammals, WIPI1 and WIPI2 also function upstream of WIPI3 and WIPI4 (26). Our data are consistent with these previous reports; *WIPI1/2* DKO cells showed an almost complete block in autophagic flux, whereas *WIPI3/4* DKO cells maintained partial autophagic activity (Fig. 1) and were able to form small autophagosomes (Fig. 3) as previously shown in neuronal cells (30), suggesting that WIPI1 and WIPI2 are important for an early step of autophagosome formation and that WIPI3 and WIPI4 are important for the elongation step. Our data showing the additive effect of WIPI2 and WIPI4 further suggest that WIPI2 and WIPI4 have distinct functions. ATG16L1 and ATG2A/B are known downstream factors of WIPI2 and WIPI4, respectively (1, 2). It was previously reported that ATG8s, which are conjugated to PE by the ATG12–ATG5–ATG16L1 complex, recruit ATG2A/B via interaction with the LIR domain in ATG2A/B (32), but the relative contribution of ATG8s and WIPI3/4 to ATG2 recruitment has been unknown. We suggest that both are redundantly important, as the LIR mutation and WIR deletion showed an additive effect (Fig. 4B and C). It should also be noted that the inhibition of Halo–GFP processing in *WIPI2* KO cells was greater than that in *ATG16L1* KO cells (Fig. 1B and C). The data suggest that WIPI2 has a yet unknown fundamental function beyond the recruitment of ATG16L1 (Fig. 8). A previous report has shown the interaction between WIPI2 and ATG2A/B (28), and we also detected a possible interaction between WIPI2B and ATG2A/B (Fig. 3C). The mutations of ATG2A, which reduce the interactions with GABARAP and WIPI3/4 and suppress the recruitment of ATG2A to phagophores, did not impair the interaction between ATG2A and WIPI2B (Fig. S4C). Therefore, the interaction between ATG2 and WIPI2 may be important for a potential mechanism other than the recruitment of ATG2. This hypothesis may be consistent with our finding that WIPI2 and WIPI4 function in autophagy through different mechanisms (Fig. 2).

BPAN patients exhibit only neuronal symptoms (33–35), and *Wipi3/4* DKO mice also demonstrate p62 accumulation only in the brain (31), leading to the hypothesis that the autophagic functions of WIPI3 and WIPI4 are specific to neuronal tissues. In this study, we revealed WIPI4 functions in autophagy, even in non-neuronal cells. Therefore, the tissue specificity of BPAN and related mouse models may be explained by the vulnerability of neuronal tissues to mild autophagy defects rather than tissue-specific functions, as seen in other autophagy-related diseases (63–67). Another possibility is that the relative importance of WIPI4 and ATG8s in recruiting ATG2A/B may differ between neuronal and non-neuronal tissues.

Halo-based reporter assays revealed various effects of BPAN-related missense mutations in WIPI4 on their autophagic activity. We found a possible association between the magnitude of autophagy defect and the severity of neurodevelopmental symptoms, which is consistent with the previous findings that neurodevelopmental defects are core symptoms in autophagy-related diseases with mutations in *EPG5*, *ATG5*, and *ATG7* (63–67). In contrast, we detected no correlation between the magnitude of autophagy defect and the age of the onset of neurodegeneration symptoms; some mutations found in BPAN patients with iron deposition and neurodegeneration symptoms showed almost normal autophagic activity. These findings suggest that the development of iron deposition and neurodegeneration in BPAN is attributable to unknown non-autophagic functions of WIPI4, for example, in retrograde transport from endosomes to the Golgi apparatus, as was also suggested for its most closely related yeast PROPPIN, Hsv2 (Fig. 8) (19). Elucidating the non-autophagic functions of WIPI4 is necessary to understand the pathogenesis of BPAN and develop therapeutics for the prevention of neurodegeneration in the disease.

## Materials and Methods

### Cell lines

HEK293T cells and HeLa cells were incubated in Dulbecco’s modified Eagle’s medium (DMEM; D6546, Sigma-Aldrich, St. Louis, MO) supplemented with 10% fetal bovine serum (FBS, 173012, Sigma-Aldrich) and 2 mM L-glutamine (25030-081, Gibco, Waltham, MA) at 37℃ in a 5% CO_2_ incubator. For starvation, after being washed twice with phosphate-buffered saline (PBS), cells were cultured in amino acid-free DMEM (048-33575, Fujifilm Wako Pure Chemical Corporation, Osaka, Japan) without serum (starvation medium) for the time indicated for each experiment. For bafilomycin A_1_ treatment, cells were incubated in a starvation medium with 100 nM bafilomycin A_1_ (B1793, Sigma-Aldrich) for the time indicated for each experiment.

### Plasmids

DNA fragments encoding WIPI1 (NM_017983.7), WIPI2 (NM_016003.4), WIPI3 (NM_019613.4), WIPI4 (NM_001029896.2), and their variants were inserted into the retroviral plasmids pMRXIP (68), pMRXIB, and pMRXIH together with 3×FLAG tag or HaloTag7 (N2701, Promega, Madison, WI). DNA fragments encoding ATG2A (NM_015104.3) and its variants were inserted into pMRXIP (68) together with monomeric ultrastable green fluorescent protein (muGFP). DNA fragments encoding ATG2A (NM_015104.3) and its variants were also inserted into pMRXNo. The pMRXIB, pMRXIH, and pMRXNo plasmids were constructed by replacing a puromycin-resistant gene cassette with a blasticidin-resistant gene cassette, a hygromycin B-resistant gene cassette, and an internal ribosomal entry site driven SNAP-tag (N9181S, New England BioLabs, Ipswich, MA), respectively. Mutated or truncated constructs were generated by PCR-mediated site-directed mutagenesis. The pMRXIP-GFP-LC3-RFP, pMRXIB-Halo-mGFP, and pMRXNo-Halo-ratLC3B plasmids were constructed as described previously (41, 42).

For the generation of KO cell lines, guide RNAs (gRNAs) were cloned into pSpCas9(BB)-2A-GFP (PX458; a gift from Dr. F. Zhang, Broad Institute of Massachusetts Institute of Technology; #48138, Addgene, Watertown, MA).

### Antibodies and reagents

For immunoblotting, the following antibodies were used: mouse monoclonal anti-HSP90 (61049, BD Transduction Lab, San Jose, CA) and anti-WIPI2 (MABC91, Sigma-Aldrich) antibodies and rabbit polyclonal anti-p62 (PM045, MBL, Tokyo, Japan), anti-ATG2A (PD041, MBL), anti-ATG2B (15131-1-AP, Proteintech, Rosemont, IL), anti-ATG16L1 (M150-3, MBL), and anti-WIPI4 (19194-1-AP, Proteintech) antibodies. Rabbit polyclonal anti-LC3 antibody was described previously (69). Horseradish peroxidase (HRP)-conjugated anti-mouse immunoglobulin G (111-035-003, Jackson ImmunoResearch Laboratories, West Grove, PA) and anti-rabbit immunoglobulin G (111-035-144, Jackson ImmunoResearch Laboratories) antibodies were used as secondary antibodies. As described above, 100 nM bafilomycin A_1_ (B1793, Sigma-Aldrich) was applied. Tetramethylrhodamine (TMR)-conjugated HaloTag ligand (G8251, Promega) and SF650-conjugated HaloTag ligand (A308-02, Goryo Chemical, Tokyo, Japan) were applied at concentrations of 100 nM and 200 nM, respectively. LipidTOX Red (H34476, Thermo Fisher Scientific, Waltham, MA) was used for the observation of lipid droplets.

### The generation of KO cell lines

The clustered regularly interspaced short palindromic repeats (CRISPR) gRNA sequences were as follows: human ATG2A, 5′-TTCCAGGTGGATGTCTCGCA-3′; human ATG2B, 5′-ATGGACTCCGAAAACGGCCA-3′; human ATG16L1, 5′-TGAATTACACAAGAAACGTG-3′; human WIPI1, 5′-CCTTATGGACAAGATGTTGC-3′; human WIPI2, 5′-GACGATGGCCACTAGGCTGC-3′; human WIPI3, 5′-GGTTGAGTGCAATGCAGCTC-3′; human WIPI4, 5′-AGACTCCCACACTTGTGTCCC-3. HEK293T cells were transfected with PX458 vector with the gRNA. After 48 h, GFP-positive cells were isolated into single clones using a cell sorter (MoFlo Astrios EQ; Beckman Coulter, Brea, CA, USA). For the confirmation of each gene KO, genomic DNAs spanning the target sequences were amplified by polymerase chain reaction (PCR), and PCR amplicons were cloned into the plasmid of the Zero Blunt TOPO PCR cloning kit (450031, Thermo Fisher Scientific) and sequenced. The PCR primers used are as follows: human ATG2A-fw, 5′-TCAGTAGGCCTCGCCCTTTGC-3′; human ATG2A-rv, 5′-AGATTCCCCTGGCCTCTCTAC-3′; human ATG2B-fw, 5′-GCCTCCTCTCGCGCTCTCTTGCACTCT-3′; human ATG2B-rv, 5′-ACACTCTAACACATTTCTCTGAGCCGT-3′; human ATG16L1-fw, 5′-TGGCTTTGTGAACATGTTTCT-3′; human ATG16L1-rv, 5′-TTTTGATGCCACATATGGCTA-3′; human WIPI1-fw, 5′-GGAAAGGACTCTGAAAGTGCTCCC-3′; human WIPI1-rv, 5′-AATCTGTACCCTAGGGAGAGGGTG-3′; human WIPI2-fw, 5′-TGTCTTTCGTGTGAATGCTCGT-3′; human WIPI2-rv, 5′-TCGTGAATGGAATGGGCTGAG-3′; human WIPI3-fw, 5′-ACCAAGAAGGGGAGACGCTTTGAG-3′; human WIPI3-rv, 5′-ACATAACTCACCACGGTGAGACCCA-3′; human WIPI4-fw, 5′-GGGGAAAGAGGCAGGGAGATG-3′; human WIPI4-rv, 5′-TGAATGGAGCAGACGAGGTG-3′.

### Preparation of retrovirus for stable expression

For the preparation of retrovirus, HEK293T cells were transfected with a retroviral vector together with pCG-VSV-G and pCG-gag-pol (gifts from Dr. T. Yasui), using Lipofectamine 2000 (11668019, Thermo Fisher Scientific). Two days after transfection, the supernatant was passed through a 0.45-μm syringe filter unit (SLHV033RB, Merck Millipore, Billerica, MA) and collected. Then, the retrovirus was applied to cells, and stable transformants were selected by application of 2 μg/mL puromycin (P8833, Sigma-Aldrich), 5 μg/mL blasticidin (022-18713, Wako), and 100 μg/mL hygromycin B (10687-010, Thermo Fisher Scientific). For the generation of *ATG2A/B* DKO HEK293T cells stably expressing ATG2A or its variants, *ATG2A/B* DKO HEK293T cells were infected with the retrovirus carrying ATG2A or ATG2A variants and the internal ribosomal entry site drive SNAP-tag. Stable transformants were labeled with 100 nM SNAP-Cell 647-SiR ligand (New England Biolabs, S9102S). The fluorescence signals from SNAP-Cell 647-SiR were detected, and cells with the same fluorescence intensities were collected by a cell sorter (MoFlo AstriosEQ; Beckman Coulter).

### Protein extraction and immunoblotting

Cells were lysed with 50 mM Tris-HCl, pH 7.5, 150 mM NaCl, 1 mM MgCl_2_, 0.2% *n*-dodecyl-β-D-maltoside, and Complete EDTA-free protease inhibitor cocktail (05056489001, Roche, Basel, Switzerland) for 10 min on ice and then treated with 0.1% benzonase (9025-65-4, Sigma-Aldrich) for 10 min on ice. The protein concentrations in the whole cell lysates were measured by the BCA method. The whole cell lysates were mixed with 6× sample buffer (46.7 mM Tris-HCl, pH 6.8, 5% glycerol, 1.67% SDS, 1% β-mercaptoethanol, and 0.02% bromophenol blue) and heated to 95℃ for 5 min. The protein concentrations were adjusted with 1×sample buffer. Samples were subjected to sodium dodecyl sulfate polyacrylamide gel electrophoresis (SDS-PAGE) and transferred to a polyvinylidene difluoride (PVDF) membrane (IPVH00010, Millipore) using the Trans-Blot Turbo Transfer System (BioRad, Hercules, CA). Immunoblotting analysis was performed with the antibodies indicated in the figures. The signals were visualized with Immobilon Western Chemiluminescent HRP Substrate (P90715, Millipore) and were detected on a Fusion SOLO.7S.EDGE (Vilber-Lourmat, Collégien, France). Contrast and brightness were adjusted and quantified using the Fiji image processing package (70).

### Coimmunoprecipitation

Cells were transfected with the indicated constructs using Lipofectamine 2000 (11668019, Thermo Fisher Scientific). Two days later, cells were lysed with 50 mM Tris-HCl, pH 7.5, 150 mM NaCl, 1 mM MgCl_2_, 0.5% NP-40, 0.1% benzonase (9025-65-4, Sigma-Aldrich), and Complete EDTA-free protease inhibitor cocktail (05056489001; Roche). Then, samples were subjected to immunoprecipitation using anti-FLAG M2 affinity gel (A2220, Sigma-Aldrich). The beads were washed three times in 50 mM Tris-HCl, pH 7.5, 150 mM NaCl, 1 mM MgCl_2_, 0.5% NP-40, and Complete EDTA-free protease inhibitor cocktail (05056489001; Roche), and bound proteins were eluted with 1× sample buffer at 95℃ for 5 min. Subsequently, samples were subjected to SDS-PAGE and immunoblotting analysis.

### Halo–GFP/LC3 processing assay

Cells were incubated with 100 nM TMR-conjugated HaloTag ligand (G8251, Promega) for 30 min. After being washed twice with PBS, cells were cultured in the starvation medium for the time indicated. Then, cells were harvested, and proteins were obtained. Proteins were separated by SDS-PAGE, and the gel was immediately visualized for in-gel fluorescence from TMR with a Fusion SOLO.7S.EDGE (Vilber-Lourmat). For the calculation of Halo–LC3 processing, the images of gels were captured with an exposure time of 30 sec. The Halo–LC3 processing rate was calculated as the intensity of the free Halo band divided by the sum of the intensities of the free Halo and unprocessed Halo–LC3 bands. In the Halo–GFP processing assay, Halo processing is usually so low that the intensity of free Halo and unprocessed Halo–GFP bands cannot be visualized in the same image because the appropriate exposure time for each is different. Therefore, the images of gels were obtained twice with exposure times of 2 s and 10 min. The Halo–GFP processing rate was calculated as the intensity of the free Halo band in the 10-min exposed image divided by the intensity of the unprocessed Halo–GFP band in the 2-s exposed image (total Halo–GFP [processed + unprocessed] can be approximated as the intensity of unprocessed Halo–GFP in a 2-s exposed image, in which processed Halo can be hardly observed).

### Flow cytometry

After a 3-min treatment with trypsin-EDTA (25300062, Gibco), cells were collected in ice-cold PBS. After washing, cells were analyzed on a cell analyzer (Cytoflex, Beckman Coulter) equipped with 488 nm, 561 nm, and 639 nm lasers. Data were processed with Kaluza analysis software (Beckman Coulter). For sorting, cells were collected in Hanks’ balanced salt solution with 20% FBS, 50 units/mL penicillin, and 50 μg/mL streptomycin (15070-063, Gibco), and then, cells were sorted by a cell sorter (MoFlo AstriosEQ; Beckman Coulter)

### Fluorescence imaging

HeLa cells expressing muGFP-ATG2A and WIPI2B-HaloTag7 were seeded onto glass-bottom dishes (3911-035, IWAKI) 48 h before observation. Cells were cultured in starvation medium with 200 nM SF650-conjugated HaloTag ligand (A308-02, Goryo Chemical) and LipidTOX Red (H34476, Thermo Fisher Scientific) for the time indicated. Then, cells were observed using a confocal laser microscope (FV3000, Olympus, Tokyo, Japan) equipped with a 60× oil immersion objective lens (1.42 NA, Olympus). Obtained images were processed using the Fiji image processing package (70). Each cell was isolated in a single image and processed individually. After being convolved with the Gaussian Blur filter in Fiji (sigma= 2), images were processed with the Top Hat filter (radius = 10). A threshold was determined for each channel and applied to all images in each channel to binarize images. Binarized images were processed by the Watershed filter, and then puncta were identified and counted by the Analyze Particles tool. Colocalized puncta were visualized by applying the Multiply function in Fiji’s Image Calculator to two binarized images of interest and then counted using Analyze Particles again.

### 3D correlative light and electron microscopy

To observe and measure autophagosomes, 3D correlative light and electron microscopy (3D-CLEM) analysis was performed as described previously (46). Briefly, gridded coverslip-bottom dishes (TCI-3922-035R-1CS, a custom-made product based on 3922-035, with cover glass attached in the opposite direction, IWAKI, Sizuoka, Japan) were coated with carbon by a vacuum evaporator (IB-29510VET, JEOL, Tokyo, Japan) and treated with 10 μg/mL poly-L-lysine for 15 min, followed by UV irradiation. Cells were seeded onto the custom-made glass-base dishes 48 h prior to the experiment. After incubation in the starvation medium for 1 h, cells were fixed with 2% paraformaldehyde (26126-54, Nacalai Tesque, Kyoto, Japan) and 0.5% glutaraldehyde (G018/1, TAAB) in 0.1 M PB, pH7.4 for 1 h at 4℃. Then, cells were observed using a confocal laser microscope (FV3000, Olympus) equipped with a 60× oil immersion objective lens (1.42 NA, Olympus). After obtaining fluorescence images, the cells were embedded in epoxy resin for observation with an electron microscope as described previously. After cells were embedded and removal of the coverslip, the resin block was trimmed into a 150 μm × 150 μm square, which included the area observed by the confocal laser microscope (FV3000, Olympus). Then, the block was cut to a 25-nm thickness to create a ribbon of 50–200 serial sections using a ultramicrotome (EM UC7, Leica, Wetzlar, Germany) equipped with a diamond knife (Ultra JUMBO, 45 degrees, DiATOME, Hatfield, PA). Those sections were observed by a scanning electron microscope (JSM7900, JEOL) with the assistance of Array Tomography Supporter software (System in Frontier, Tokyo, Japan). The images were stacked in order using Stacker NEO TEMography.com 3.3.4.0 software (System in Frontier). The alignment of the images obtained with the confocal laser microscope and the electron microscope was done using Fiji software. The maximum cross-sectional areas of autophagosomes were also measured using Fiji software. The observation of autophagosomes, the quantification of autophagosome sizes, and the statistical analysis were performed in a double-blinded manner.

### Statistical analysis

Statistical analyses were performed with GraphPad Prism 8 software (GraphPad Software, Boston, MA). The statistical method used for each experiment is described in each figure legend.

## Acknowledgements

We thank Tatsushi Toda for his mentorship and encouragement, Yoko Ishida and Keiko Igarashi for technical assistance with the 3D-CLEM experiment, Shoji Yamaoka (Tokyo Medical and Dental University, Tokyo, Japan) for pMRXIP, and Teruhito Yasui (National Institutes of Biomedical Innovation, Health and Nutrition (NIBIOHN), Osaka, Japan) for pCG-VSV-G and pCG-gag-pol.

## Funding

This work was supported by the Exploratory Research for Advanced Technology (ERATO) research funding program of the Japan Science and Technology Agency (JST) (JPMJER1702 to NM) and a Grant-in-Aid for Specially Promoted Research from the Japan Society for the Promotion of Science (JSPS) (22H04919 to NM).

**Figure S1. CRISPR/Cas9-mediated gene knockout was confirmed by genomic DNA sequencing and immunoblotting.**

(A–I) Genomic DNA sequences of targeted regions are shown. Insertions are highlighted in red, and deletions are indicated by dashes. Frameshift mutations were confirmed in *WIPI1* KO, *WIPI2* KO, *WIPI3* KO, *WIPI4* KO, *WIPI1/2* DKO, *WIPI3/4* DKO, and *WIPI1–4* QKO HEK293T cells. In *ATG16L1* KO HEK293T cells, a frameshift mutation was detected in one allele and an in-frame deletion involving a splice site was detected in the other. In *ATG2A/B* DKO HEK293T cells, frameshift mutations were detected in both alleles of *ATG2A* genes. In *ATG2B* genes, a frameshift mutation was detected in one allele, and an in-frame mutation disrupting an initiator codon was detected in the other. (J) Immunoblotting of total cell lysates of the indicated HEK293T cell lines is shown. No commercially available antibody can detect endogenous WIPI1 and WIPI3.

**Figure S2. The LC3 and p62 turnover and GFP–LC3–RFP reporter methods did not reveal a decrease in autophagic flux in *WIPI3/4* DKO HEK293T cells.**

(A) WT and *WIPI3/4* DKO HEK293T cells were incubated for 4 h in regular or starvation medium with or without 100 nM bafilomycin A_1_.

(B) and (C) The band intensities of LC3-II and p62 normalized with those of HSP90 were quantified. Data from three experiments are plotted. Data were statistically analyzed by Welch’s *t*-test.

(D) Schematic representations of the measurement of autophagic flux using the GFP–LC3–RFP reporter. GFP–LC3–RFP is cleaved by ATG4 family proteins to produce equimolar amounts of GFP–LC3 and RFP. GFP–LC3 is efficiently engulfed and degraded by autophagosomes, while RFP remains in the cytosol and serves as an internal control. The reduction of the GFP:RFP ratio reflects the autophagic flux.

(E) WT and *WIPI3/4* DKO HEK293T cells stably expressing GFP–LC3–RFP were incubated for 12 h in regular or starvation medium. The fluorescence intensity of GFP and RFP were determined by flow cytometry. Geometric means of the fluorescence intensity of GFP over that of RFP (GFP:RFP ratio) were calculated and normalized with the values in regular medium. Data from three experiments were statistically analyzed by Welch’s *t*-test.

**Figure S3. The WIPI4 N15A/D17A mutation abolishes the interaction between WIPI4 and ATG2A/B.**

Cell lysates from HEK293T cells transiently expressing 3×FLAG-tagged WPI4 or its mutants were subjected to immunoprecipitation with anti-FLAG antibody and immunoblotting with antibodies against FLAG and endogenous ATG2A and ATG2B.

**Figure S4. The formation of autophagosomes is not impaired in *WIPI3/4* DKO cells.**

(A) WT and *WIPI3/4* DKO HEK293T cells expressing GFP–LC3 were incubated in starvation medium for 1 h; scale bar, 5 μm.

(B) The number of autophagosomes was counted in three WT and *WIPI3/4* DKO cells. Solid bars indicate the means, and dots indicated the data from three different cells. Data were statistically analyzed using the Mann–Whitney *U*-test.

**Figure S5. The LC3-interacting region and WIPI-interacting region in ATG2A are important for binding with ATG8 and WIPI4, respectively.**

Either 3×FLAG-tagged GABARAP (A), WIPI4 (B), or WIPI2B (C) were transiently co-expressed with muGFP, muGFP–ATG2A, or the indicated ATG2A mutants in HEK293T cells. Cell lysates were subjected to immunoprecipitation with anti-FLAG antibody and immunoblotting with antibodies against FLAG and GFP.

## Abbreviations

ATG: autophagy-related
BPAN: β-propeller associated neurodegeneration
ER: endoplasmic reticulum
DKO: double knockout
KO: knockout
PI3P: phosphatidylinositol 3-phosphate
PROPPINs: β-propellers that bind polyphosphoinositides
QKO: quadruple knockout
SENDA: static encephalopathy of childhood with neurodegeneration in adulthood
WIPI: WD-repeat protein interacting with phosphoinositides

## References

1. Yamamoto, H., Zhang, S., Mizushima, N. (2023) Autophagy genes in biology and disease. Nat. Rev. Genet., 10.1038/s41576-022-00562-w.

2. Nakatogawa, H. (2020) Mechanisms governing autophagosome biogenesis. Nat. Rev. Mol. Cell. Biol., 21, 439–458.

3. Klionsky, D. J., Cregg, J. M., Dunn, W. A., Jr, Emr, S. D., Sakai, Y., Sandoval, I. V., Sibirny, A., Subramani, S., Thumm, M., Veenhuis, M. et al. (2003) A unified nomenclature for yeast autophagy-related genes. Dev. Cell, 5, 539–545.

4. Kotani, T., Kirisako, H., Koizumi, M., Ohsumi, Y., & Nakatogawa, H. (2018). The Atg2-Atg18 complex tethers pre-autophagosomal membranes to the endoplasmic reticulum for autophagosome formation. Proc. Natl. Acad. Sci. U. S. A., 115, 10363–10368.

5. Osawa, T., Kotani, T., Kawaoka, T., Hirata, E., Suzuki, K., Nakatogawa, H., Ohsumi, Y., & Noda, N. N. (2019). Atg2 mediates direct lipid transfer between membranes for autophagosome formation. Nat. Struct. Mol. Biol., 26, 281–288.

6. Valverde, D. P., Yu, S., Boggavarapu, V., Kumar, N., Lees, J. A., Walz, T., Reinisch, K. M., & Melia, T. J. (2019). ATG2 transports lipids to promote autophagosome biogenesis. J. Cell. Biol., 218, 1787–1798.

7. Mizushima, N., Noda, T., Yoshimori, T., Tanaka, Y., Ishii, T., George, M. D., Klionsky, D. J., Ohsumi, M., & Ohsumi, Y. (1998). A protein conjugation system essential for autophagy. Nature, 395, 395–398.

8. Mizushima, N., Kuma, A., Kobayashi, Y., Yamamoto, A., Matsubae, M., Takao, T., Natsume, T., Ohsumi, Y., & Yoshimori, T. (2003). Mouse Apg16L, a novel WD-repeat protein, targets to the autophagic isolation membrane with the Apg12-Apg5 conjugate. J. Cell. Sci., 116, 1679–1688.

9. Mizushima, N., Sugita, H., Yoshimori, T., & Ohsumi, Y. (1998). A new protein conjugation system in human. The counterpart of the yeast Apg12p conjugation system essential for autophagy. J. Biol. Chem., 273, 33889–33892.

10. Proikas-Cezanne, T., Waddell, S., Gaugel, A., Frickey, T., Lupas, A., & Nordheim, A. (2004). WIPI-1alpha (WIPI49), a member of the novel 7-bladed WIPI protein family, is aberrantly expressed in human cancer and is linked to starvation-induced autophagy. Oncogene, 23, 9314–9325.

11. Polson, H. E., de Lartigue, J., Rigden, D. J., Reedijk, M., Urbé, S., Clague, M. J., & Tooze, S. A. (2010) Mammalian Atg18 (WIPI2) localizes to omegasome-anchored phagophores and positively regulates LC3 lipidation. Autophagy, 6, 506–522.

12. Barth, H., Meiling-Wesse, K., Epple, U. D., & Thumm, M. (2001) Autophagy and the cytoplasm to vacuole targeting pathway both require Aut10p. FEBS lett., 508, 23–28.

13. Guan, J., Stromhaug, P. E., George, M. D., Habibzadegah-Tari, P., Bevan, A., Dunn, W. A., Jr, & Klionsky, D. J. (2001) Cvt18/Gsa12 is required for cytoplasm-to-vacuole transport, pexophagy, and autophagy in Saccharomyces cerevisiae and Pichia pastoris. Mol. Biol. Cell, 12, 3821–3838.

14. Obara, K., Sekito, T., Niimi, K., & Ohsumi, Y. (2008) The Atg18-Atg2 complex is recruited to autophagic membranes via phosphatidylinositol 3-phosphate and exerts an essential function. J. Biol. Chem., 283, 23972–23980.

15. Watanabe, Y., Kobayashi, T., Yamamoto, H., Hoshida, H., Akada, R., Inagaki, F., Ohsumi, Y., & Noda, N. N. (2012) Structure-based analyses reveal distinct binding sites for Atg2 and phosphoinositides in Atg18. J. Biol. Chem., 287, 31681–31690.

16. Meiling-Wesse, K., Barth, H., Voss, C., Eskelinen, E. L., Epple, U. D., & Thumm, M. (2004) Atg21 is required for effective recruitment of Atg8 to the preautophagosomal structure during the Cvt pathway. J. Biol. Chem., 279, 37741–37750.

17. Strømhaug, P. E., Reggiori, F., Guan, J., Wang, C. W., & Klionsky, D. J. (2004) Atg21 is a phosphoinositide binding protein required for efficient lipidation and localization of Atg8 during uptake of aminopeptidase I by selective autophagy. Mol. Biol. Cell, 15, 3553–3566.

18. Juris, L., Montino, M., Rube, P., Schlotterhose, P., Thumm, M., & Krick, R. (2015) PI3P binding by Atg21 organises Atg8 lipidation. EMBO. J., 34, 955–973.

19. Bueno-Arribas, M., Blanca, I., Cruz-Cuevas, C., Escalante, R., Navas, M. A., & Vincent, O. (2021) A conserved ATG2 binding site in WIPI4 and yeast Hsv2 is disrupted by mutations causing β-propeller protein-associated neurodegeneration. Hum. Mol. Genet., 31, 111–121.

20. Dove, S. K., Piper, R. C., McEwen, R. K., Yu, J. W., King, M. C., Hughes, D. C., Thuring, J., Holmes, A. B., Cooke, F. T., Michell, R. H. et al. (2004) Svp1p defines a family of phosphatidylinositol 3,5-bisphosphate effectors. EMBO. J., 23, 1922–1933.

21. Gopaldass, N., Fauvet, B., Lashuel, H., Roux, A., & Mayer, A. (2017) Membrane scission driven by the PROPPIN Atg18. EMBO. J., 36, 3274–3291.

22. Courtellemont, T., De Leo, M. G., Gopaldass, N., & Mayer, A. (2022) CROP: a retromer-PROPPIN complex mediating membrane fission in the endo-lysosomal system. EMBO. J., 41, e109646.

23. Marquardt, L., Taylor, M., Kramer, F., Schmitt, K., Braus, G. H., Valerius, O., & Thumm, M. (2023) Vacuole fragmentation depends on a novel Atg18-containing retromer-complex. Autophagy, 19, 278–295.

24. Lu, Q., Yang, P., Huang, X., Hu, W., Guo, B., Wu, F., Lin, L., Kovács, A. L., Yu, L., & Zhang, H. (2011) The WD40 repeat PtdIns(3)P-binding protein EPG-6 regulates progression of omegasomes to autophagosomes. Dev. Cell, 21, 343–357.

25. Dooley, H. C., Razi, M., Polson, H. E., Girardin, S. E., Wilson, M. I., & Tooze, S. A. (2014) WIPI2 links LC3 conjugation with PI3P, autophagosome formation, and pathogen clearance by recruiting Atg12-5-16L1. Mol. Cell, 55, 238–252.

26. Bakula, D., Müller, A. J., Zuleger, T., Takacs, Z., Franz-Wachtel, M., Thost, A. K., Brigger, D., Tschan, M. P., Frickey, T., Robenek, H. et al. (2017) WIPI3 and WIPI4 β-propellers are scaffolds for LKB1-AMPK-TSC signalling circuits in the control of autophagy. Nat. Commun., 8, 15637.

27. Chowdhury, S., Otomo, C., Leitner, A., Ohashi, K., Aebersold, R., Lander, G. C., & Otomo, T. (2018) Insights into autophagosome biogenesis from structural and biochemical analyses of the ATG2A-WIPI4 complex. Proc. Natl. Acad. Sci. U. S. A., 115, E9792–E9801.

28. Zheng, J. X., Li, Y., Ding, Y. H., Liu, J. J., Zhang, M. J., Dong, M. Q., Wang, H. W., & Yu, L. (2017) Architecture of the ATG2B-WDR45 complex and an aromatic Y/HF motif crucial for complex formation. Autophagy, 13, 1870–1883.

29. Ren, J., Liang, R., Wang, W., Zhang, D., Yu, L., & Feng, W. (2020) Multi-site-mediated entwining of the linear WIR-motif around WIPI β-propellers for autophagy. Nat. Commun., 11, 2702.

30. Ji, C., Zhao, H., Chen, D., Zhang, H., & Zhao, Y. G. (2021) β-propeller proteins WDR45 and WDR45B regulate autophagosome maturation into autolysosomes in neural cells. Curr. Biol., 31, 1666–1677

31. Ji, C., Zhao, H., Li, D., Sun, H., Hao, J., Chen, R., Wang, X., Zhang, H., & Zhao, Y. G. (2020) Role of *Wdr45b* in maintaining neural autophagy and cognitive function. Autophagy, 16, 615–625.

32. Bozic, M., van den Bekerom, L., Milne, B. A., Goodman, N., Roberston, L., Prescott, A. R., Macartney, T. J., Dawe, N., & McEwan, D. G. (2020) A conserved ATG2-GABARAP family interaction is critical for phagophore formation. EMBO. Rep., 21, e48412.

33. Saitsu, H., Nishimura, T., Muramatsu, K., Kodera, H., Kumada, S., Sugai, K., Kasai-Yoshida, E., Sawaura, N., Nishida, H., Hoshino, A. et al. (2013) De novo mutations in the autophagy gene WDR45 cause static encephalopathy of childhood with neurodegeneration in adulthood. Nat. Genet., 45, 445–449.

34. Haack, T. B., Hogarth, P., Kruer, M. C., Gregory, A., Wieland, T., Schwarzmayr, T., Graf, E., Sanford, L., Meyer, E., Kara, E. et al. (2012) Exome sequencing reveals de novo WDR45 mutations causing a phenotypically distinct, X-linked dominant form of NBIA. Am. J. Hum. Genet., 91, 1144–1149.

35. Hayflick, S. J., Kruer, M. C., Gregory, A., Haack, T. B., Kurian, M. A., Houlden, H. H., Anderson, J., Boddaert, N., Sanford, L., Harik, S. I. et al. (2013) β-Propeller protein-associated neurodegeneration: a new X-linked dominant disorder with brain iron accumulation. Brain, 136, 1708–1717.

36. Najmabadi, H., Hu, H., Garshasbi, M., Zemojtel, T., Abedini, S. S., Chen, W., Hosseini, M., Behjati, F., Haas, S., Jamali, P. et al. (2011) Deep sequencing reveals 50 novel genes for recessive cognitive disorders. Nature, 478, 57–63.

37. Suleiman, J., Allingham-Hawkins, D., Hashem, M., Shamseldin, H. E., Alkuraya, F. S., & El-Hattab, A. W. (2018) WDR45B-related intellectual disability, spastic quadriplegia, epilepsy, and cerebral hypoplasia: A consistent neurodevelopmental syndrome. Clin. Genet., 93, 360–364.

38. Almannai, M., Marafi, D., Abdel-Salam, G. M. H., Zaki, M. S., Duan, R., Calame, D., Herman, I., Levesque, F., Elbendary, H. M., Hegazy, I. et al. (2022) El-Hattab-Alkuraya syndrome caused by biallelic WDR45B pathogenic variants: Further delineation of the phenotype and genotype. Clin. Genet., 101, 530–540.

39. Jelani, M., Dooley, H. C., Gubas, A., Mohamoud, H. S. A., Khan, M. T. M., Ali, Z., Kang, C., Rahim, F., Jan, A., Vadgama, N. et al. (2019) A mutation in the major autophagy gene, WIPI2, associated with global developmental abnormalities. Brain, 142, 1242–1254.

40. Maroofian, R., Gubas, A., Kaiyrzhanov, R., Scala, M., Hundallah, K., Severino, M., Abdel-Hamid, M. S., Rosenfeld, J. A., Ebrahimi-Fakhari, D., Ali, Z. et al. (2021) Homozygous missense *WIPI2* variants cause a congenital disorder of autophagy with neurodevelopmental impairments of variable clinical severity and disease course. Brain. Commun., 3, fcab183.

41. Yim, W. W., Yamamoto, H., & Mizushima, N. (2022) A pulse-chasable reporter processing assay for mammalian autophagic flux with HaloTag. Elife, 11, e78923.

42. Kaizuka, T., Morishita, H., Hama, Y., Tsukamoto, S., Matsui, T., Toyota, Y., Kodama, A., Ishihara, T., Mizushima, T., & Mizushima, N. (2016) An Autophagic Flux Probe that Releases an Internal Control. Mol. Cell, 64, 835–849.

43. Mauthe, M., Jacob, A., Freiberger, S., Hentschel, K., Stierhof, Y. D., Codogno, P., & Proikas-Cezanne, T. (2011) Resveratrol-mediated autophagy requires WIPI-1-regulated LC3 lipidation in the absence of induced phagophore formation. Autophagy, 7, 1448–1461.

44. Baskaran, S., Ragusa, M. J., Boura, E., & Hurley, J. H. (2012) Two-site recognition of phosphatidylinositol 3-phosphate by PROPPINs in autophagy. Mol. Cell, 47, 339–348.

45. Liang, R., Ren, J., Zhang, Y., & Feng, W. (2019) Structural Conservation of the Two Phosphoinositide-Binding Sites in WIPI Proteins. J. Mol. Biol., 431, 1494–1505.

46. Takahashi, S., Saito, C., Koyama-Honda, I., & Mizushima, N. (2022) Quantitative 3D correlative light and electron microscopy of organelle association during autophagy. Cell. Struct. Funct., 47, 89–99.

47. Velikkakath, A. K., Nishimura, T., Oita, E., Ishihara, N., & Mizushima, N. (2012). Mammalian Atg2 proteins are essential for autophagosome formation and important for regulation of size and distribution of lipid droplets. Mol. Biol. Cell, 23, 896–909.

48. Tamura, N., Nishimura, T., Sakamaki, Y., Koyama-Honda, I., Yamamoto, H., & Mizushima, N. (2017) Differential requirement for ATG2A domains for localization to autophagic membranes and lipid droplets. FEBS Lett., 591, 3819–3830.

49. Kobayashi, T., Suzuki, K., & Ohsumi, Y. (2012) Autophagosome formation can be achieved in the absence of Atg18 by expressing engineered PAS-targeted Atg2. FEBS Lett., 586, 2473–2478.

50. Zarate, Y. A., Jones, J. R., Jones, M. A., Millan, F., Juusola, J., Vertino-Bell, A., Schaefer, G. B., & Kruer, M. C. (2016) Lessons from a pair of siblings with BPAN. Eur. J. Hum. Genet., 24, 1080–1083.

51. Khoury, J., Kotagal, P., & Moosa, A. N. V. (2019) Epileptic encephalopathy and brain iron accumulation due to WDR45 mutation. Seizure, 71, 245–246.

52. Long, M., Abdeen, N., Geraghty, M. T., Hogarth, P., Hayflick, S., & Venkateswaran, S. (2015) Novel WDR45 Mutation and Pathognomonic BPAN Imaging in a Young Female With Mild Cognitive Delay. Pediatrics, 136, e714–e717.

53. Chard, M., Appendino, J. P., Bello-Espinosa, L. E., Curtis, C., Rho, J. M., Wei, X. C., & Al-Hertani, W. (2019) Single-center experience with Beta-propeller protein-associated neurodegeneration (BPAN); expanding the phenotypic spectrum. Mol. Genet. Metab. Rep., 20, 100483.

54. Chen, H., Qian, Y., Yu, S., Xiao, D., Guo, X., Wang, Q., Hao, L., Yan, K., Lu, Y., Dong, X. et al. (2019) Early onset developmental delay and epilepsy in pediatric patients with WDR45 variants. Eur. J. Med. Genet., 62, 149–160.

55. Russo, C., Ardissone, A., Freri, E., Gasperini, S., Moscatelli, M., Zorzi, G., Panteghini, C., Castellotti, B., Garavaglia, B., Nardocci, N. et al. (2018) Substantia Nigra Swelling and Dentate Nucleus T2 Hyperintensity May Be Early Magnetic Resonance Imaging Signs of β-Propeller Protein-Associated Neurodegeneration. Mov. Disord. Clin. Pract., 6, 51–56.

56. Kim, M. K., Kim, N. Y., Hong, S., Ma, H. I., & Kim, Y. J. (2017) Presynaptic Dopaminergic Degeneration in a Patient with Beta-Propeller Protein-Associated Neurodegeneration Documented by Dopamine Transporter Positron Emission Tomography Images: A Case Report. J. Mov. Disord., 10, 161–163.

57. Carvill, G. L., Liu, A., Mandelstam, S., Schneider, A., Lacroix, A., Zemel, M., McMahon, J. M., Bello-Espinosa, L., Mackay, M., Wallace, G. et al. (2018) Severe infantile onset developmental and epileptic encephalopathy caused by mutations in autophagy gene WDR45. Epilepsia, 59, e5–e13.

58. Tschentscher, A., Dekomien, G., Ross, S., Cremer, K., Kukuk, G. M., Epplen, J. T., & Hoffjan, S. (2015) Analysis of the C19orf12 and WDR45 genes in patients with neurodegeneration with brain iron accumulation. J. Neurol. Sci., 349, 105–109.

59. Nishioka, K., Oyama, G., Yoshino, H., Li, Y., Matsushima, T., Takeuchi, C., Mochizuki, Y., Mori-Yoshimura, M., Murata, M., Yamasita, C. et al. (2015) High frequency of beta-propeller protein-associated neurodegeneration (BPAN) among patients with intellectual disability and young-onset parkinsonism. Neurobiol. Aging, 36, 2004.e9–2004.e15.

60. Verhoeven, W. M., Egger, J. I., Koolen, D. A., Yntema, H., Olgiati, S., Breedveld, G. J., Bonifati, V., & van de Warrenburg, B. P. (2014) Beta-propeller protein-associated neurodegeneration (BPAN), a rare form of NBIA: novel mutations and neuropsychiatric phenotype in three adult patients. Parkinsonism Relat. Disord., 20, 332–336.

61. De Leo, M. G., Berger, P., & Mayer, A. (2021) WIPI1 promotes fission of endosomal transport carriers and formation of autophagosomes through distinct mechanisms. Autophagy, 17, 3644–3670.

62. Tsukida, K., Muramatsu, S. I., Osaka, H., Yamagata, T., & Muramatsu, K. (2022) *WDR45* variants cause ferrous iron loss due to impaired ferritinophagy associated with nuclear receptor coactivator 4 and WD repeat domain phosphoinositide interacting protein 4 reduction. Brain Commun., 4, fcac304.

63. Dionisi Vici, C., Sabetta, G., Gambarara, M., Vigevano, F., Bertini, E., Boldrini, R., Parisi, S. G., Quinti, I., Aiuti, F., & Fiorilli, M. (1988) Agenesis of the corpus callosum, combined immunodeficiency, bilateral cataract, and hypopigmentation in two brothers. Am. J. Med. Genet., 29, 1–8.

64. Cullup, T., Kho, A. L., Dionisi-Vici, C., Brandmeier, B., Smith, F., Urry, Z., Simpson, M. A., Yau, S., Bertini, E., McClelland, V. et al. (2013) Recessive mutations in EPG5 cause Vici syndrome, a multisystem disorder with defective autophagy. Nat Genet., 45, 83–87.

65. al Shahwan, S. A., Bruyn, G. W., & al Deeb, S. M. (1995) Non-progressive familial congenital cerebellar hypoplasia. J. Neurol. Sci., 128, 71–77.

66. Kim, M., Sandford, E., Gatica, D., Qiu, Y., Liu, X., Zheng, Y., Schulman, B. A., Xu, J., Semple, I., Ro, S. H. et al. (2016) Mutation in ATG5 reduces autophagy and leads to ataxia with developmental delay. Elife, 5, e12245.

67. Collier, J. J., Guissart, C., Oláhová, M., Sasorith, S., Piron-Prunier, F., Suomi, F., Zhang, D., Martinez-Lopez, N., Leboucq, N., Bahr, A. et al. (2021) Developmental Consequences of Defective ATG7-Mediated Autophagy in Humans. N. Engl. J. Med., 384, 2406–2417.

68. Kitamura, T., Koshino, Y., Shibata, F., Oki, T., Nakajima, H., Nosaka, T., & Kumagai, H. (2003) Retrovirus-mediated gene transfer and expression cloning: powerful tools in functional genomics. Exp. Hematol., 31, 1007–1014.

69. Kabeya, Y., Mizushima, N., Ueno, T., Yamamoto, A., Kirisako, T., Noda, T., Kominami, E., Ohsumi, Y., & Yoshimori, T. (2000) LC3, a mammalian homologue of yeast Apg8p, is localized in autophagosome membranes after processing. EMBO. J., 19, 5720–5728.

70. Schindelin, J., Arganda-Carreras, I., Frise, E., Kaynig, V., Longair, M., Pietzsch, T., Preibisch, S., Rueden, C., Saalfeld, S., Schmid, B. et al. (2012) Fiji: an open-source platform for biological-image analysis. Nat. Methods., 9, 676–682.

