## Supplemental figures for "Comprehensive analysis of autophagic functions of WIPI family proteins and their implications for the pathogenesis of β-propeller associated neurodegeneration"

**A *WIPI1* KO**

| | |
|---|---|
| *WIPI1 WT allele* | CTCCAGCAACATCTTGTCCATAAGGCTG |
| *WIPI1* KO allele-1 | CTCCAGCAAACATCTTGTCCATAAGGCT |
| *WIPI1* KO allele-2 | CTCCAGCAA———————————TG |

**B *WIPI2* KO**

| | |
|---|---|
| *WIPI2* WT allele | TGTTCTCCAGCAGCCTAGTGGCCATC |
| *WIPI2* KO allele-1 | TGTTCTCCAGC–GCCTAGTGGCCATC |
| *WIPI2* KO allele-2 | TGTTCTCCAGCAGGCCTAGTGGCCAT |

**C *WIPI3* KO**

| | |
|---|---|
| *WIPI3* WT allele | GAGGGTGTCCTGAGCTGCATTGCACTC |
| *WIPI3* KO allele-1 | GAGGGTGTCCTGAG——GCATTGCACTC |
| *WIPI3* KO allele-2 | GAGGGTGTC————CTGCATTGCACTC |

**D *WIPI4* KO**

| | |
|---|---|
| *WIPI4* WT allele | GTTCCCGGGACACAAGTGTGGGAGT |
| *WIPI4* KO allele-1 | GTTCCCGG—————————GGGAGT |
| *WIPI4* KO allele-2 | GTTCCCGG–ACACAAGTGTGGGAGT |

**E *WIPI1/2* DKO**

| | | | |
|---|---|---|---|
| *WIPI1* WT allele | CTCCAGCAACATCTTGTCCATAAGGCTG | *WIPI2* WT allele | TGTTCTCCAGCAGCCTAGTGGCCATC |
| *WIPI1* KO allele-1 | CTCCAGCAAACATCTTGTCCATAAGGCT | *WIPI2* KO allele-1 | G———————————GCCTAGTGGCCAT |
| *WIPI1* KO allele-2 | CTCCAGCAA———————————TG | *WIPI2* KO allele-2 | TGTTCTCCAGCAGGCCTAGTGGCCAT |

**F *WIPI3/4* DKO**

| | | | |
|---|---|---|---|
| *WIPI3* WT allele | TTTCCGGGCACGCACACGGGCCATGTG | *WIPI4* WT allele | TGTTCCCGGGACACAAGTGTGGGAGT |
| *WIPI3* KO allele-1 | TTT—————CGCACACGGGCCATGTG | *WIPI4* KO allele-1 | TGT——————————————GGGAGT |
| *WIPI3* KO allele-2 | TTT——————————————TGTG | *WIPI4* KO allele-2 | TGTTCCCGG—ACACAAGTGTGGGAGT |

**G *WIPI1-4* QKO**

| | | | |
|---|---|---|---|
| *WIPI1* WT allele | CTCCAGCAACATCTTGTCCATAAGGCTG | *WIPI2* WT allele | ATTGTTCTCCAGCAGCCTAGTGGCCAT |
| *WIPI1* KO allele-1 | CTCCAGCAAACATCTTGTCCATAAGGCT | *WIPI2* KO allele-1 | ATTGTTC—————GCCTAGTGGCCAT |
| *WIPI1* KO allele-2 | A—(409bp insertion)—ACATCTTGTCCAT | *WIPI2* KO allele-2 | AT——————————GCACAAAGTGGC |
| *WIPI3* WT allele | TTTCCGGGCACGCACACGGGCCATGTG | *WIPI4* WT allele | AGTGTTCCCGGGACACAAGTGTGGGAG |
| *WIPI3* KO allele-1 | TT——————————————TG | *WIPI4* KO allele-1 | AGTGT———————————GGGAG |
| *WIPI3* KO allele-2 | TTTCCCGGGGCACGCACACGGGCCATG | *WIPI4* KO allele-2 | AGTGTTCCCGG–ACACAAGTGTGGGAG |

**H *ATG16L1* KO**

| | |
|---|---|
| *ATG16L1* WT allele | AGAAACGTGGGGAGGTAAAGCT |
| *ATG16L1* KO allele-1 | AG——————————————CT |
| *ATG16L1* KO allele-2 | AGAAACCGTGGGGAGGTAAAGC |

**I *ATG2A/B* DKO**

| | |
|---|---|
| *ATG2A* WT allele | GCAGCGTTGCCCTGCGAG |
| *ATG2A* KO allele-1 | GC————————GAG |
| *ATG2A* KO allele-2 | GCAGCGTTGCCCTGCGGA |
| *ATG2B* WT allele | CCGTTTTCGGAGTCCATCAAGAAGAGGGCCTGCCGGTACCTCCTGC |
| *ATG2B* KO allele-1 | ————(78bp deletion)————GGTCTGCGGGCATAGTGAC |
| *ATG2B* KO allele-2 | CCCGTTTTCGGAGTCCATCAAGAAGAGGGCCTGCCGGTACCTCCTG |

**J**

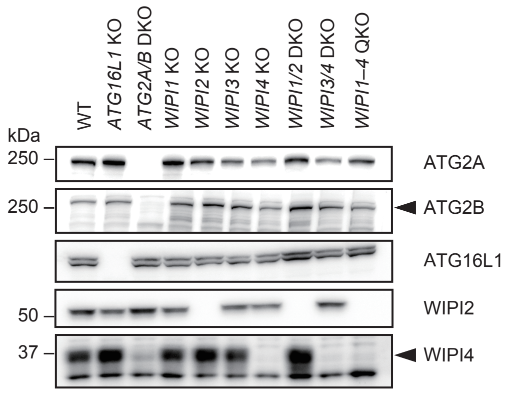


**Figure S1. CRISPR/Cas9-mediated gene knockout was confirmed by genomic DNA sequencing and immunoblotting.**

(A-I) Genomic DNA sequences of targeted regions are shown. Insertions are highlighted in red and deletions are indicated by dashes. Frameshift mutations were confirmed in *WIPI1* KO, *WIPI2* KO, *WIPI3* KO, *WIPI4* KO, *WIPI1/2* DKO, *WIPI3/4* DKO, and *WIPI1–4* QKO HEK293T cells. In *ATG16L1* KO HEK293T cells, a frameshift mutation was detected in one allele and an in-frame deletion involving a splice site was detected in the other. In *ATG2A/B* DKO HEK293T cells, frameshift mutations were detected in both alleles of *ATG2A* genes. In *ATG2B* genes, a frameshift mutation was detected in one allele, and an in-frame mutation disrupting an initiator codon was detected in the other.

(J) Immunoblotting of total cell lysates of the indicated HEK293T cell lines is shown. No commercially available antibody can detect endogenous WIPI1 and WIPI3.

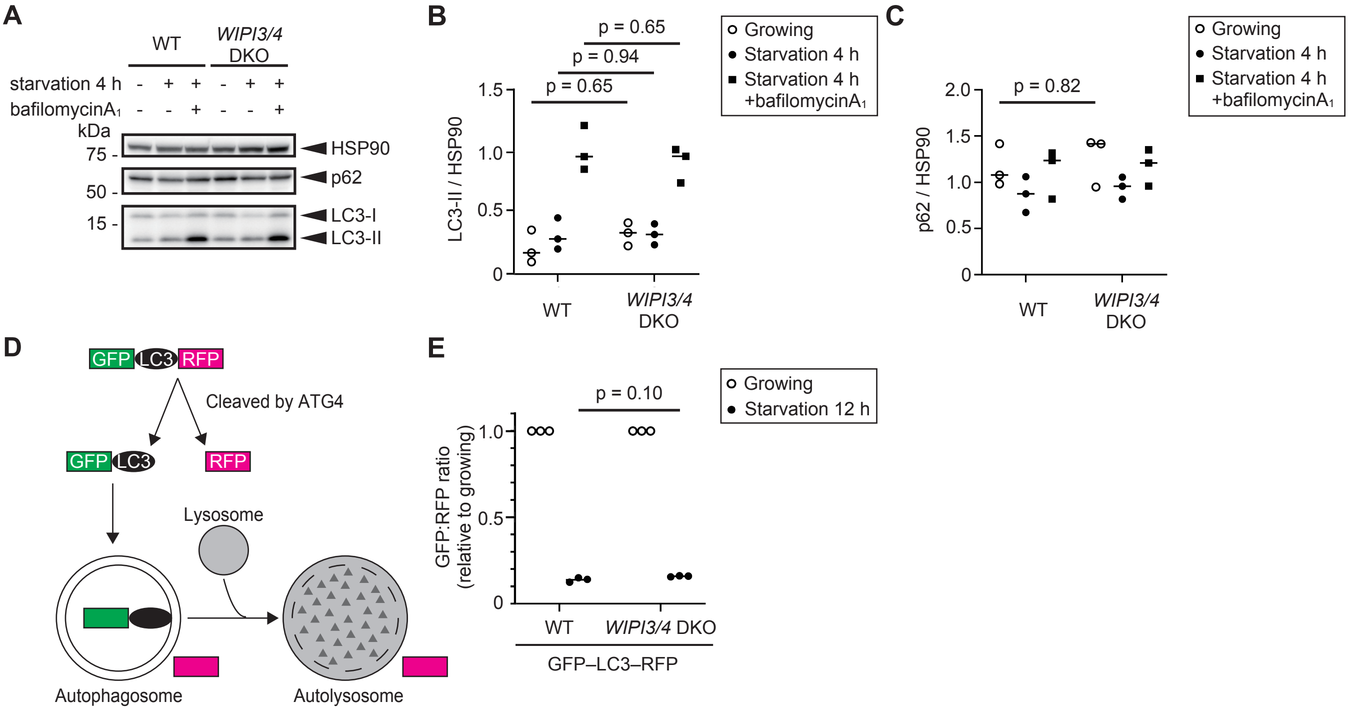


**Figure S2. The LC3 and p62 turnover and GFP–LC3–RFP reporter methods did not reveal a decrease in autophagic flux in *WIPI3/4* DKO HEK293T cells.**

(A) WT and *WIPI3/4* DKO HEK293T cells were incubated for 4 h in regular or starvation medium with or without 100 nM bafilomycin A1.

(B) and (C) The band intensities of LC3-II and p62 normalized with those of HSP90 were quantified. Data from three experiments are plotted. Data were statistically analyzed by by Holm-Šídák test (B) and Welch' s *t*-test (C)..

(D) Schematic representations of the measurement of autophagic flux using the GFP–LC3–RFP reporter. GFP–LC3–RFP is cleaved by ATG4 family proteins to produce equimolar amount of GFP–LC3 and RFP. GFP–LC3 is efficiently engulfed and degraded by autophagosomes, while RFP remains in the cytosol and serves as an internal control. The reduction of the GFP:RFP ratio reflects the autophagic flux.

(E) WT and *WIPI3/4* DKO HEK293T cells stably expressing GFP–LC3–RFP were incubated for 12 h in regular or starvation medium. The fluorescence intensity of GFP and RFP were determined by flow cytometry. Geometric means of the fluorescence intensity of GFP over that of RFP (GFP:RFP ratio) were calculated and normalized with the values in regular medium. Data from three experiments were statistically analyzed by Welch' s *t*-test.

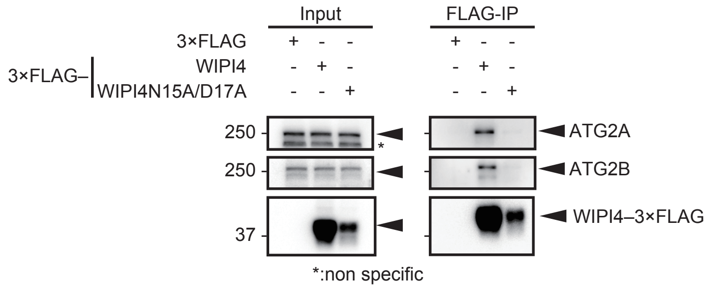


**Figure S3. WIPI4 N15A/D17A mutation abolishes the interaction between WIPI4 and ATG2A/B.**

Cell lysates from HEK293T cells transiently expressing 3×FLAG-tagged WPI4 or its mutants were subjected to immunoprecipitation with anti-FLAG antibody and immunoblotting with antibodies against FLAG and endogenous ATG2A and ATG2B.

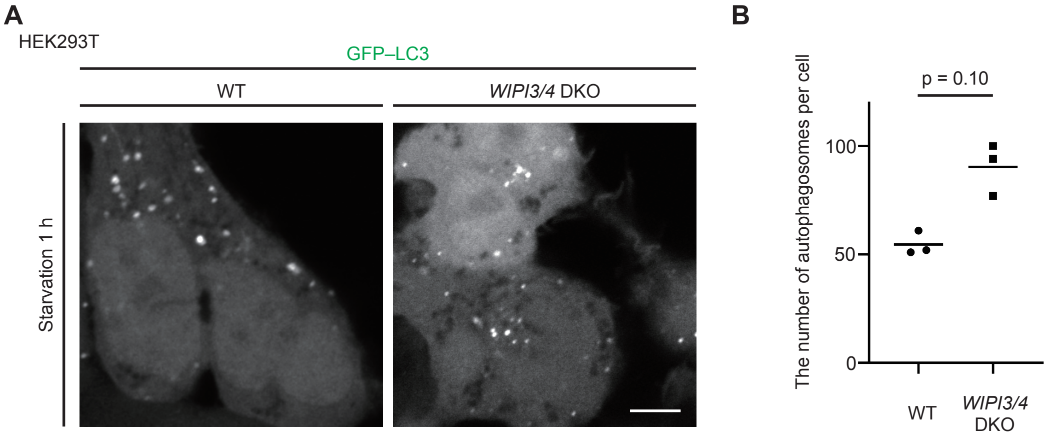


**Figure S4. The formation of autophagosomes is not impaired in *WIPI3/4* DKO cells.**

(A) WT and *WIPI3/4* DKO HEK293T cells expressing GFP–LC3 were incubated in starvation medium for 1 h; Scale bar, 5 μm.

(B) The number of autophagosomes was counted in three WT and *WIPI3/4* DKO cells. Solid bars indicate the means and dots indicated the data from three different cells. Data were statistically analyzed using the Mann–Whitney *U*-test.

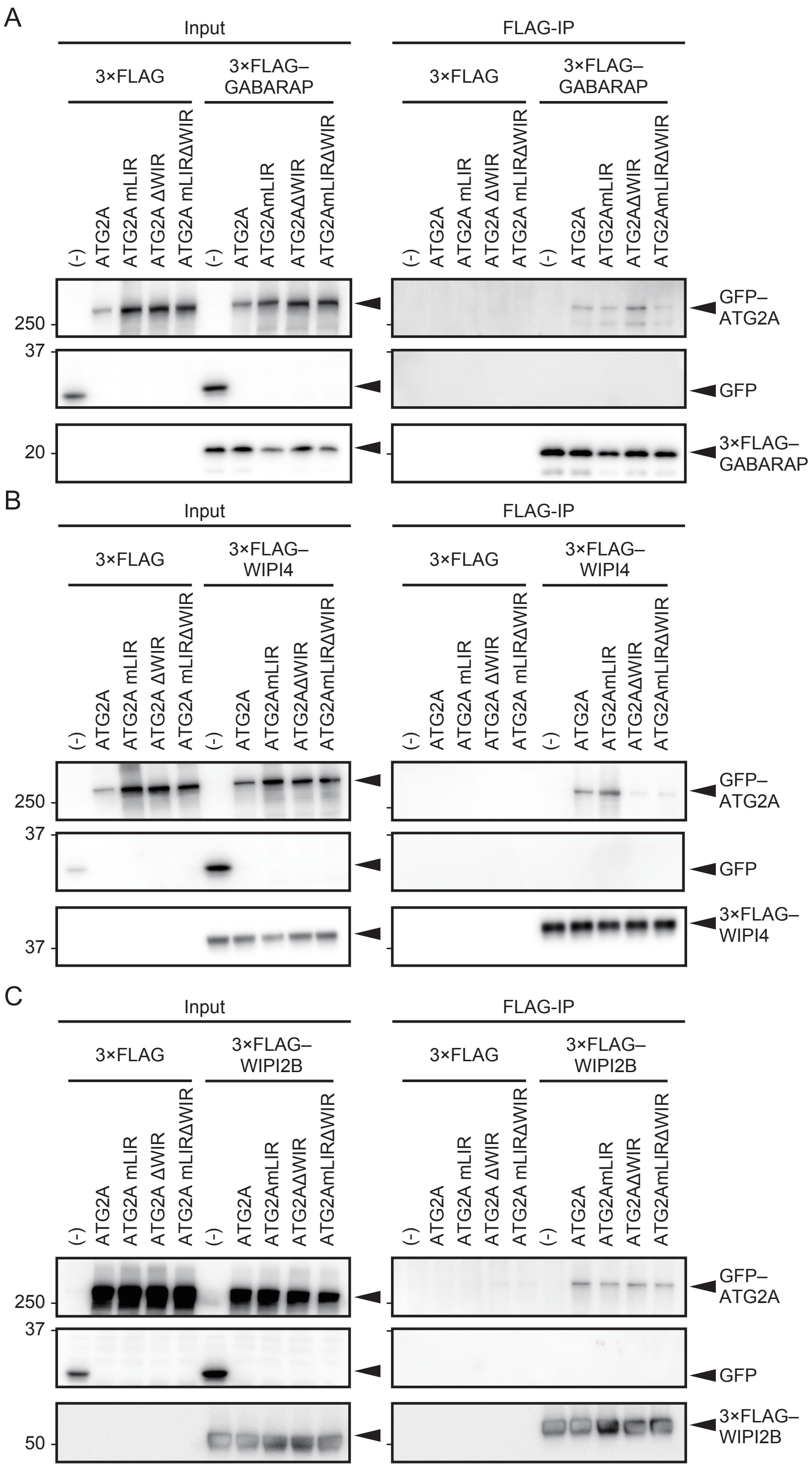


**Figure S5. The LC3-interacting region and WIPI-interacting region in ATG2A are important for binding with ATG8 and WIPI4, respectively.**

Either 3×FLAG-tagged GABARAP (A), WIPI4 (B), or WIPI2B (C) were transiently co-expressed with muGFP, muGFP–ATG2A, or the indicated ATG2A mutants in HEK293T cells. Cell lysates were subjected to immunoprecipitation with anti-FLAG antibody and immunoblotting with antibodies against FLAG and GFP.
